# Immortalization and Characterization of Schwann Cell Lines Derived from NF1 Associated Cutaneous Neurofibromas

**DOI:** 10.1101/2025.03.19.643702

**Authors:** Hua Li, Alexander Pemov, Robert Allaway, David F. Muir, Lung-Ji Chang, Jineta Banerjee, Alexandra J. Scott, Jaime M.W. Nagy, Jian Liu, Meritxell Carrió, Helena Mazuelas, Anthony Yachnis, Sang Y. Lee, Xiaochun Zhang, Yang Lyu, Douglas R. Stewart, Angela Hirbe, Jaishri O. Blakeley, Eduard Serra, Deeann Wallis, Margaret R. Wallace

## Abstract

Neurofibromatosis type 1 (NF1) is an autosomal dominant condition in which patients are heterozygous for a disruptive pathogenic variant in the *NF1* gene. The most characteristic feature of the condition NF1 is the neurofibroma, a benign, multi-cellular tumor which initiates when a cell of the Schwann cell lineage gains a somatic pathogenic variant of the other *NF1* allele. Neurofibromas developing at nerve termini in the skin are termed “cutaneous” neurofibromas (cNFs), while those developing within larger nerves are termed “plexiform.” Most patients develop cNFs beginning in late childhood or early adulthood, continuing throughout life at variable rates. Some patients may develop only a few cNFs, while others suffer from thousands. There are no reliably effective physical or pharmaceutical therapies besides surgical removal. Although these are not life-threatening, they are disfiguring and can interfere with normal life functions. To provide a resource for research, we developed short-term cNF Schwann cell cultures from NF1 patients, from which we subsequently established the first semi-immortalized cNF cell lines through transduction with wild-type human telomerase reverse transcriptase (*hTERT*) and murine cyclin-dependent kinase 4 (*mCdk4*) genes. Here we present molecular, cellular, and functional characterization of these cell lines, which will be of utility for investigating and developing *NF1* cNF therapies.

---

Neurofibromatosis type 1 (NF1) is an autosomal dominant condition with birth incidence estimated at 1/2700.^1^ NF1 patients are germline heterozygous for a *NF1* gene pathogenic variant (formerly termed “mutation”), with half of these inherited from an affected parent, and half arising through new mutation at the gamete level, although a small minority of cases are mosaic (post-zygotic mutation).^2^ Although a number of different clinical features characterize NF1, it was named for the presence of neurofibromas, benign Schwann cell (SC) tumors that can arise anywhere on the peripheral nervous system. These masses additionally contain axons, fibroblasts, perineurial cells, and often some mast cells and CD34+ cells.^3–5^ Neurofibromas that develop in the skin, arising at nerve termini, are referred to as cutaneous neurofibromas (cNFs), and rarely become larger than an inch in diameter. Typically, cNFs become evident in adolescence and increase throughout life in an unpredictable fashion, although in some cases, hormonal changes can stimulate their growth.^6^ A few rare *NF1* pathogenic variants are associated with lack of cNFs, but this is highly unusual; most patients develop multiple cNFs over their lifespan, ranging from dozens to thousands.^1^ Therapies to prevent or shrink cNFs are in early development, and excision by surgery, shaving, or laser are tedious and may only have a temporary effect. cNFs impact quality of life through disfigurement, itch, pain, and/or functional impingement.^7^ In contrast to cNFs, neurofibromas developing within large nerves are termed “plexiform.” Plexiform neurofibromas (pNFs) may grow very large with substantial morbidity and potential mortality, especially if they transform to a malignant peripheral nerve sheath tumor (MPNST).^8^ cNFs are generally not associated with malignant transformation, however, there are rare instances of diffuse cNF that share cNF and plexiform features.^9,10^

*NF1* encodes neurofibromin, a 250-kD tumor suppressor protein that is a negative regulator of RAS signaling (encoded by middle eighth of the 8454-bp 57-exon coding region (for isoform 2, encoding 2818-amino acid minimal protein, ENST00000356175)), as well as other less-characterized functions.^11^ All data to date shows that NF1 neurofibroma development requires a somatic *NF1* pathogenic variant of the other allele in a SC (called “two-hit,” *NF1* -/-), consistent with the Knudson tumor two-hit phenomenon.^12–14^ Thousands of different intragenic germline and somatic *NF1* pathogenic variants have been reported in manuscripts and databases. It is thought that neurofibromas arise when a *NF1* two-hit SC begins proliferating in response to wound-like environment signals (even without an actual wound), regardless of contact with an axon, and stimulates production of stroma as well.^15^ Genome-wide studies have not identified highly consistent, driver somatic variants at other loci in neurofibroma SCs, except where the somatic pathogenic variant is a deletion spanning *NF1* and surrounding genes.^16–18^

Cell cultures consisting of tumor cells can be critical research tools for investigating the biology of, and testing therapeutics for, tumors. Cultivation of SCs from tissue is not always successful, and requires a substrate and growth factor supplementation.^13,19^ Wild-type human SC cultures rarely divide beyond 6-8 passages, and often have residual fibroblasts. From cNFs, derived cultures may also contain both heterozygous and two-hit SC, although conditions can be used to enrich for one type or the other.^13^ Several years ago, to address these issues and provide improved resources for NF1 translational research, we discovered that human SC cultures (including wild type, and *NF1* +/- and *NF1* -/- from pNFs) can be “immortalized” by transduction with expression constructs for both human telomerase reverse transcriptase (*hTERT*) and murine cyclin-dependent kinase 4 (*mCdk4*) genes.^20^ These cell lines are able to divide many passages beyond the original culture (in some cases more than 80 passages), and most no longer require a substrate or growth factor for propagation. These cell lines have been widely used in NF1 research aimed at understanding and treating pNFs, as well as in other SC-related conditions and studies. However, given the difference in natural history and molecular profiles between pNFs and cNFs, it is likely that cNF-derived cell lines will have unique properties that will inform translational research focused on these tumors. We report here the production of immortalized cNF SC lines, as well as their molecular and functional characterization.

## Materials and Methods

### Subjects

Eight individuals planning to undergo surgery to remove cNFs gave informed consent to donate their tumor samples and a blood sample to research in compliance with relevant laws and approval from the University of Florida (UF) IRB, protocols 41-1992 (1992-2012) and 2017-00320 (2013-current). A total of 14 cNF samples were collected through this process (Table 1, Supplemental Table 1). All subjects met diagnostic criteria for NF1,^21^ except for one (a young child lost to follow up, cell line icNF97.5, no matched blood sample). Three additional cNF primary SC cultures used for immortalization (18cNF, 21cNF, 28cNF) were derived from anonymized samples under approval of the Clinical Research Ethics Committee of Germans Trias i Pujol Hospital, Badalona (Barcelona), Spain; no demographic or clinical information was available except a diagnosis of NF1.

### Schwann cell (SC) primary cultures

Primary cultures of cNF-derived SCs were established from surgically-removed neurofibromas as previously described,^20,22^ in DMEM medium with 10% fetal bovine serum and antibiotics, using a substrate (laminin or collagen I) and neuregulin supplementation (glial growth factor 2). For three cNF (18cNF, 21cNF, 28cNF) initial SCs were isolated as described elsewhere.^13^

### hTERT and mCdk4 lentiviral vector transduction

The vector pTYcPPT, containing the elongation factor 1 alpha promoter,^23^ was used to separately clone *hTERT* (full length human telomerase reverse transcriptase cDNA) and *mCdk4* (murine *Cdk4* full-length cDNA). Based on previous experience creating pNF immortalized cells,^20^ cNF cultures (10,000 cells) were transduced using both lentiviruses simultaneously with 10 multiplicity of infection (MOI) for both viruses, for 12-18 hours with 10 µg/ml polybrene. Media was then changed, and cells were maintained on 0.1% collagen 1-treated plates (Sigma, C8919) in high glucose DMEM with L-glutamine, 10% fetal bovine serum, 1X Antibiotic-Antimycotic (Fisher 15240062), 5 µg/ml Plasmocin (Invivogen, ANT-MPP), and 50 ng/µl neuregulin (R&D Systems 396-HB-050). These cultures were expanded and passaged; when cells continued to proliferate beyond the passage at which the original primary culture senesced, the cell line was labeled “immortalized” and was designated with an “i” in front of the culture name.

### hTERT and mCdk4 reverse transcription (RT)-PCR

This experiment was used to confirm transgene expression, using the primers and conditions described previously.^20^ Since telomerase is not detectable by RT-PCR in unmodified cultured SC,^20^ detection of *hTERT* was used to verify *hTERT* transgene expression. The *Cdk4* RT-PCR primers are specific for the murine cDNA, and thus detection of expression confirmed presence and activity of that transgene.

### Cell line authentication and characterization of proliferation, apoptosis, karyotype, and S100B expression

The cell lines had variable capacity to divide before senescing, ranging from p.22 to p42 or further passages. They were first screened for the ability to survive and proliferate without a substrate and/or neuregulin. The cell lines were also evaluated for apoptotic rate (TUNEL staining), proliferation rate (BrdU incorporation), and S100B expression, using previously published methods.^22^ Cell authentication using STR genotyping utilized the Promega GenePrint 10 system, comparing DNA from primary cultures to immortalized cell lines, at the University of Florida’s Interdisciplinary Center for Biotechnology Research. Routine G-banded karyotypes were obtained for 12 cell lines by established cytogenetics laboratories. Cell lines were regularly tested for mycoplasma contamination.

### Subcloning a heterozygous line from icNF98.4c

In an effort to create isogenic cell line pairs for several tumors, we successfully subcloned a heterozygous version of icNF98.4c cells (row 16, Table 1). Cells were diluted and placed into a 96-well plate at an average of 1 cell/well. Forty single-cell clones were tested for lack of the somatic pathogenic mutation (deletion), that is, presence of two alleles at the site of the germline variant, and propagated. A subclone was found to be heterozygous with no evidence of the two-hit cells (not further characterized at this time).

### Western blots and analysis: neurofibromin and pERK:ERK

Cells from all lines reported in Table 1 were lysed with RIPA buffer and lysates were cleared by centrifugation at 20,000 RPM for 30 minutes at 4°C. Protein was quantitated with a Bradford assay; 50 µg of protein was loaded per well for NF1 blots and 20 µg of protein was loaded for other blots. 8% SDS-polyacrylamide gels were run at 100 V for 2 hours and transferred at 100 V for 2 hours onto a PVDF membrane. Blots (3 replicates) were probed overnight at 4°C with primary antibody washed and probed 1 hour at room temperature with secondary antibody. Primary antibodies included N-Terminal NF1 (Cell Signaling cat# D7R7D 1:1000), beta-actin (Cell Signaling cat# 3700 1:10000), p-ERK (Cell Signaling cat# 9101 1:1000), and total ERK (Cell Signaling cat# 9102 1:1000). The secondary antibody was HRP-tagged from Santa Cruz, for chemiluminescent detection using chemiluminescent substrate from BioRad (Cat. #: 170-5061; Hercules, California) per the manufacturer’s protocols. Western blot band intensities were quantitated with ImageJ software. Neurofibromin (when present) was normalized to beta actin of that sample. Comparison of neurofibromin, p-ERK, and ERK levels across blots was accomplished by normalizing to the signal from the heterozygous cell line icNF09.5.

### Histological analysis of original neurofibromas

For the neurofibromas acquired under University of Florida approval, formalin-fixed, paraffin embedded (FFPE) slide sections were obtained. For each tumor, one section each was stained with established methods at the University of Florida Molecular Pathology Core lab, for: (1) hematoxylin and eosin (H & E) to evaluate histology, cellularity, and vascularity; (2) toluidine blue to identify mast cells, and (3) CD69 immunostaining to identify macrophages. Stained slides were evaluated by a neuropathologist.

### NF1 Mutation Analyses

The search for *NF1* pathogenic variants in the primary cNF cultures and corresponding immortalized cell lines initially utilized Sanger sequencing of the *NF1* cDNA via overlapping reverse-transcription-PCR fragments, and Multiplex Ligation-Dependent Probe Amplification (MLPA) analysis of DNA to detect copy number changes (insertions/deletions) at multiple points in the *NF1* gene (P082 kit, MRC Holland). DNA was extracted with the PureGene system (Qiagen), and RNA extracted using the DirectZol kit (Zymo Research). cDNA sequencing included the immediate flanking 5’ and 3’ untranslated regions, but not the entire such regions (primers listed in Supplemental Table 2). cDNA variants were characterized at the DNA level, and vice versa. For five samples, high-throughput sequencing of the *NF1* locus was also employed, utilizing the Ion S5 system (ThermoFisher Scientific, Waltham, MA). Briefly, a targeted multiplex PCR primer panel was designed using custom Ion AmpliSeq Designer (ThermoFisher Scientific). The primer panel (n=393) covered all *NF1* exons and a limited intronic sequence adjacent to the exons, totaling 12 kb. Sample DNA (30 ng) was amplified using this custom AmpliSeq primer panel, and libraries were prepared following the manufacturer’s protocol for Ion AmpliSeq Library Kit 2.0. Individual samples were barcoded during library preparation and subsequently pooled. Automated template preparation and loading onto Ion 520 Chip was performed using Ion Chef system, and DNA was sequenced on the Ion S5 sequencer per manufacturer’s instructions. Variant calling of the germline samples was performed using The Genome Analysis Toolkit (https://gatk.broadinstitute.org) and IonTorrent Variant Caller (TVC, https://github.com/domibel/IonTorrent-VariantCaller), while somatic variant calling was performed using either MuTect (https://github.com/broadinstitute/mutect) or TVC with relaxed stringency in order to increase sensitivity. Germline variant pathogenicity classification was performed by AutoGVP.^24^ Somatic variants were queried in cBioPortal (https://www.cbioportal.org/) using OncoQuery Language (OCL) for each variant across all studies that have mutation data. Samples were filtered out that did not have *NF1* profiled in the sequencing panel.

### Exome, genome, and transcriptome sequencing of primary and immortalized cNF cells

Seven primary SC cultures and their matched immortalized cell lines were chosen for genomic analysis (Table 1; icNF97.2a, icNF97.2b, icNF98.4c, icNF98.4d, icNF00.10a, icNF04.9a, i28cNF). We did not perform genomic profiling on cell lines where NF1 diagnostic criteria was not met (icNF97.5), cell lines where metadata about the parent tumor was limited (i18cNF, i21cNF), and cell lines where a somatic *NF1* mutation was low frequency, not observed, or unclear (icNF18.1, icNF09.5, icNF93.1, icNF99.1, i18cNF). For whole exome sequencing (WES), exomes were captured by IDT exome reagent xGen Exome Research Panel V1.0 (Integrated DNA Technologies) and libraries were subsequently constructed with a Kapa Hyper amplified kit (Roche) and sequenced using a NovaSeq 6000 S4 (Illumina) with ∼200× coverage. For whole genome sequencing (WGS), libraries were constructed with Kapa Hyper PCR-free kits and sequenced using the NovaSeq 6000 S4 with 30x coverage. For RNA-seq, samples were prepared with non-stranded library prep with RiboErase for rRNA depletion. The libraries were sequenced on the NovaSeq6000 S4 with targeted coverage of 30M reads per sample.

### Bioinformatic and statistical analyses of whole exome and whole genome sequencing data

WES and WGS data processing was performed with a modified version of the Nextflow nf-core “sarek” data processing pipeline (v2.7.1).^25–28^ The pipeline was modified to add DeepVariant as an additional variant-calling method.^28^ Details about all processing steps and underlying software versions can be found in the nf-core/sarek documentation (https://nf-co.re/sarek/2.7.1).^29–36^ Briefly, the sequencing data were preprocessed using GATK best practice methods and aligned to the GRCh38 genome (GATK build). As there were no patient-matched non-tumor samples available, variant calling was performed in tumor-only mode and thus variations include both germline and somatic mutations. Whole exome variant calling was performed with FreeBayes,^37^ Mutect2, Strelka2,^38^ and DeepVariant.^28^ WGS variant calling was performed with the same variant calling tools. For downstream analysis, variants were filtered with bcftools, removing all variants that were not marked as a “PASS” by their respective variant caller, or variants that had a quality score less than 20. Variants were then annotated using Ensembl VEP v107 and simultaneously converted to Mutation Annotation Format (maf) using vcf2maf (https://github.com/mskcc/vcf2maf). The analyses presented here use only variants called by DeepVariant, however, the dataset (see Data Availability) includes the variant calls from the other aforementioned variant calling tools. In addition, copy-number alterations were determined using Control-FREEC (whole genome data only).^39^ Filtered variant (vcf) files for each cell line identified by DeepVariant were compared using the “jaccard” function from the “bedtools” software package.^40^ The Jaccard indices were calculated in a pairwise manner (*i.e.*, all cell lines compared to all other cell lines), followed by hierarchical clustering of the resulting matrix of Jaccard indices, plotting the results with the R “pheatmap” package (https://cran.r-project.org/web/packages/pheatmap/index.html). For the lollipop analysis, *NF1* variants were extracted from the annotated maf files from the WES data and plotted using the “maftools” R package.^41^ Copy-number alterations (CNAs) called by Control-FREEC were summarized by averaging the copy number values in 500 base pair windows across the genome. The Circlize package was used to generate circos plots to compare the CNAs to the gene counts.^42,43^ The genes in the gene expression data were mapped to the appropriate chromosome and genomic start location using the BiomaRt R package.^44,45^ The CNA values and gene counts were log10 transformed to make smaller changes easier to observe in resulting plots.

Presence of hTERT and mCdk4 transgene integration in the immortalized cell line genomes was confirmed by creating a custom genome (hg38 with additional sequences representing the hTERT and mCdk4 cDNA, which span two or more exons). We then aligned all of the WGS files to this custom genome, and evaluated the presence or absence of alignment of these additional sequences to the WGS data using Integrative Genomics Viewer.

### Bioinformatic and statistical analyses of gene expression

Transcriptomic data processing was performed using the nf-core “rnaseq” data processing pipeline (v3.5).^46^ Details about all processing steps performed and underlying software versions can be found in the nf-core/rnaseq documentation (https://nf-co.re/rnaseq/3.5). The pipeline was run in STAR-Salmon mode, which uses STAR for alignment (GRCh38, NCBI build), and Salmon for quantification of transcripts and genes. The pairwise Euclidean distances between gene expression data was also calculated during data processing by the nf-core/rnaseq pipeline.^47–49^ The raw transcriptomics counts of the samples generated using Salmon were first loaded into R. On detection of duplicate rows with same gene names, rows containing higher (max) counts were preserved and others were filtered out. The raw counts were then filtered to remove genes with counts lower than 10 in greater than 80% of the samples. The filtered counts were then normalized to account for differences in library preparation using the trimmed mean of M-values (TMM method).^50,51^ Principal component analysis using the normalized log transformed counts (log counts per million) was performed using the “prcomp” function of the “stats” package and visualized using “ggplot2.” The normalized counts were adjusted for heteroscedasticity using the “voom” function then analyzed by fitting a generalized linear model (limma and edgeR packages) to identify differentially expressed genes between the sample groups.^50–55^ “Pheatmap” was used for hierarchical clustering of the differentially expressed genes and visualizing the gene expression levels (log counts per million) between samples. A hypergeometric test using the “gprofiler2” package was performed on the differentially expressed gene set to explore the enrichment of pathways from Gene Ontology (GO), Kyoto Encyclopedia of Genes and Genomes (KEGG), Reactome (REAC), WikiPathways (WP), Transcription factor (TF), MicroRNA (MIRNA), Human Protein Atlas (HPA), CORUM (database of mammalian protein complexes), and Human Phenotype Ontology (HP) in the selected gene sets.^55,56^ Pathway terms with a term size less than 500 enriched in the selected gene sets were explored further.

### Data Resources

Cellosaurus^57^ entries have been created for the 7 cell lines (and matched 7 primary SC cultures) that were sequenced, containing information such as STR profiles and Research Resource Identifiers, and other information. This information has also been deposited in the NF Research Tools Database.^58^ Raw and processed data as well as analyses are available on Synapse.org and the NF Data Portal (https://doi.org/10.7303/syn11374339).^59^ The processed genomics data are available at syn29529772 (RNA-seq), syn29346753 (WES), and syn35832496 (WGS). Raw genomic datasets with specimen metadata are available at syn29390037.1 (RNA-seq), syn39613693.1 (WES), and syn39613866.1 (WGS). Analysis code is available at https://github.com/nf-osi/cnf-cell-lines. All analyses were performed in R 4.2.1. All R packages and versions have been recorded in a renv lockfile available at: https://github.com/nf-osi/cnf-cell-lines/blob/main/renv.lock.

## Results

### Cell lines

Co-transduction of 14 primary cNF SC cultures with lentiviral vectors containing wild type hTERT and mCdk4 cDNA transgenes led to creation of cell lines that passage well beyond the primary cultures, usually with some morphological changes (often to a shorter, stubbier shape than the typical spindle Schwann-cell morphology). Table 1 and Supplemental Table 1 summarizes the cell line parameters and characterizations, as well as lists a single-cell-cloned heterozygous line derived from the icNF98.4c cell line that was not separately characterized (row 16). Table 1 and Supplemental Table 1 also includes data from histopathology analysis of the corresponding primary neurofibroma tissue (where available), as well as information about retained dependence on, or preference for, a collagen I matrix and SC growth factor neuregulin. Presence and expression of transgenes in the cell lines was confirmed in the WGS, and through detection of transcripts by RT-PCR (data not shown). Figure 1 shows example photomicrographs from two immortalized cell lines (phase contrast, and immunostained with S100B, a SC marker).

**Figure 1.**
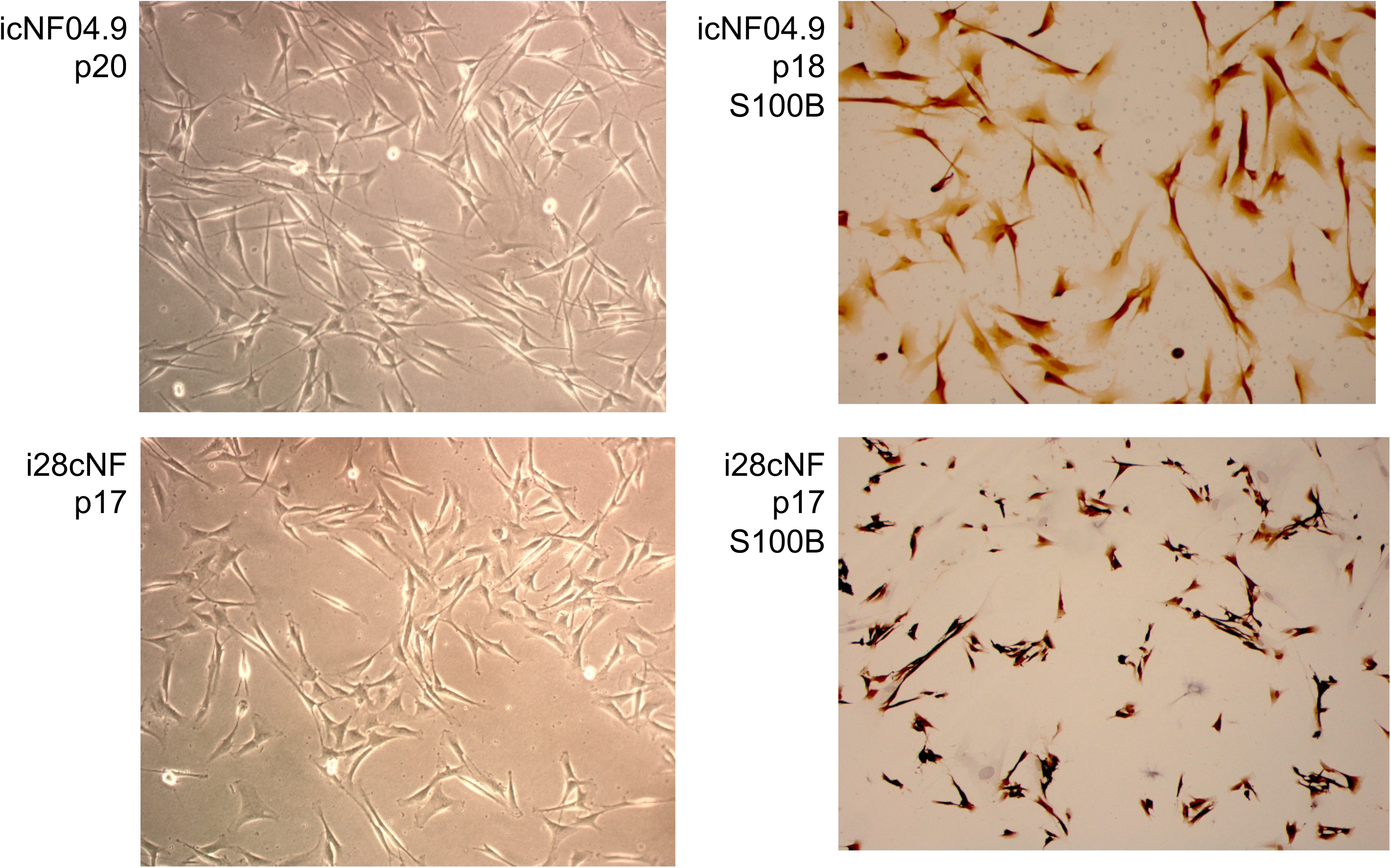
Photomicrographs of phase-contrast and S100B immunostaining of two immortalized cNF cell lines, as examples, icNF04.9 and i28cNF. The immortalized cells tend to be moderately shorter and broader than the typical long thin spindle shape of Schwann cells. However, the strong S100B staining indicates that these are in the Schwann cell lineage. A small minority of cells failed to stain with S100B, which represent other cell type(s) from the primary culture such as fibroblasts.

### NF1 variants

Table 1 also lists the *NF1* germline and somatic variant data derived from the *NF1*-focused molecular analyses. Details of germline and somatic *NF1* variants from WGS are shown in Supplemental Table 3 and Supplemental Table 4, respectively. All the germline variants were classified as pathogenic or likely pathogenic, with the exception of icNF97.5. It harbors an *NF1* synonymous substitution near the 3’ end of the coding sequence, c.8160G>C (NM_000267, isoform 2) p.Thr2720= that is listed in gnomAD 4.0 (rs777755159) but with a frequency of 0.000006816 (11/1,613,850). It has been reported twice in ClinVar in NF1 cases and is classified as “Likely Benign.” cDNA sequencing across flanking exons showed that this allele is not associated with altered mRNA splicing or quantity. If this is a *NF1* pathogenic or hypomorphic variant, it would involve a different mechanism, such as affecting translation rate due to mRNA folding or codon usage differences.^60^

The other *NF1* germline variants were nonsense or frameshift alterations (two from aberrant splicing), a missense variant, and one in-frame exon skip. Unlike the other pathogenic variants, the transcript from the p.Arg1362Ter germline pathogenic allele associated with icNF18.1a and icNF09.5 was barely detectable, suggesting a high degree of nonsense mediated decay (NMD). Interestingly, germline variant c.233del does not display NMD despite its close proximity to the 5’ end of the coding region (exon 3). It has recently been shown that missense *NF1* pathogenic variants may significantly alter neurofibromin stability and activity through affecting its dimerization, even if the new amino acid has the same general chemical properties.^61,62^

A wide range of somatic *NF1* pathogenic variants (Supplemental Table 4) was also observed. These include: splice errors, loss of heterozygosity (presumed deletion of one entire *NF1* locus, endpoints undetermined), frameshift deletions, nonsense, and one missense variant. A somatic pathogenic variant could not be identified in three cell lines (“heterozygous?” in Table 1; rows 7-9), which also had low S100B staining. In two other lines, the somatic pathogenic variant was detected at a low level (designated “mostly heterozygous” in Table 1; rows 6 and 12). In a few cases, the somatic mutation had not been detected in the primary culture analysis, but was detected in the immortalized cells, suggesting enrichment of the two-hit cells. Germline and somatic variants were scattered across the gene, and were also observed in whole exome and whole genome profiling of the cell lines (Supplemental Figures 1 and 2, Supplemental Table 5).

### Western blot analysis

Western blots of protein from the tumor-derived immortal cell lines were used to detect neurofibromin and determine the phosphorylated ERK (pERK, active form) to total ERK ratio as a downstream measure of RAS activity, reflecting neurofibromin RAS-GTPase activating function. These results vary somewhat between cell lines, consistent with different *NF1* genotypes. Representative Western blots are shown in Figure 2A, with quantitation in Figures 2B and 2C. A whole-gene deletion, nonsense or frameshift *NF1* variant are expected to result in a dramatic reduction or absence of functional neurofibromin. In cell lines heterozygous for such variants, any full-length-sized neurofibromin Western signal should represent protein encoded by the other allele. Caveats in predicting and interpreting such data, particularly from in-frame deletions or exon skips, missense pathogenic variants, and synonymous variants, include variable NMD, leaky splicing or protein stability. In addition, the presence of any “contaminating” cell may contribute to neurofibromin detection. For example, the low but detectable level of neurofibromin in i18cNF could be due to leaky splicing or presence of heterozygous cells in the culture. This latter possibility is consistent with the observation of a small percentage of cells negative for S100B in most cell lines. The icNF93.1a cell line expresses low levels of neurofibromin despite lack of a detectable somatic *NF1* variant, suggesting that the germline variant (p.Trp784Arg) may result in protein instability. In fact, this variant maps to neurofibromin’s cysteine-serine-rich domain (CSRD) which is important for several neurofibromin interactions, and some CSRD missense mutations are associated with a more severe phenotype.^61^ Similarly, icNF00.10a does not show any full-length neurofibromin, suggesting that the amino acid substitution results in protein instability. As expected, all cNF cell lines show pERK:ERK ratios elevated compared to the wild type control ipNF97.4 cells, which have very low pERK levels. icNF98.4c has the lowest pERK:ERK ratio, which is difficult to interpret given that the germline pathogenic variant results in out-of-frame skipping of exon 43, and the somatic mutation is deletion of the gene, which together predicts high pERK. icNF97.5 also has relatively low pERK:ERK, but this may reflect the unidentified somatic variant (if any) or the function of the synonymous variant. The moderate pERK:ERK ratio observed in i18cNF may suggest that the exon-skipped transcript is translated into a somewhat stable and functional protein.

**Figure 2.**
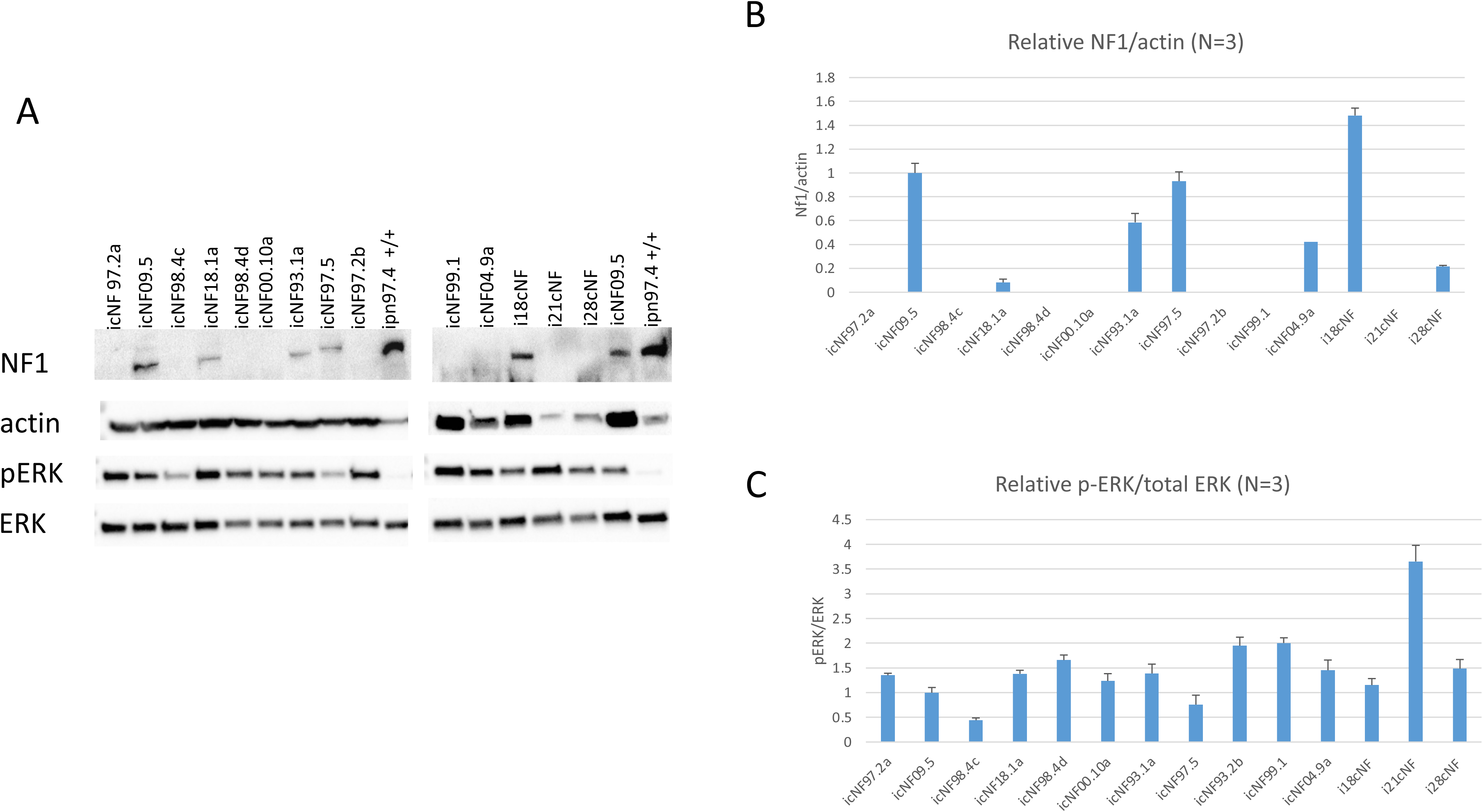
*NF1* and ERK Western blot analysis of cNF cell line protein extracts. Panel A shows cropped results from the same blot hybridized for neurofibromin (NF1, 250 kD) and subsequently beta actin (42 kD), phosphorylated ERK (pERK, 44 kD, active form) and total ERK protein (ERK, 42 kD). Immortalized wild type human Schwann cells were used as positive control for neurofibromin (ipn97.4). Panel B is a bar graph of abundance of neurofibromin (NF1) from three Western blots, relative to beta actin for each cell line (mean with 1 standard deviation), normalized to score from icNF09.5 heterozygous cells. Seven cell lines had undetectable neurofibromin. Panel C shows quantitation of the pERK:ERK signal from the three Westerns. Larger scores represent a greater proportion of ERK in the active form, which is expected with decreased neurofibromin activity.

### Bioinformatic analysis of matched primary and immortalized cell line exomes and genomes

Jaccard indices (to evaluate genomic similarity) were calculated from exome sequencing data of the immortalized cNF lines and the unmodified primary cell cultures. The primary cell cultures, when compared to their matched immortalized counterparts, had higher Jaccard indices than with unrelated cultures/lines (Figure 3 and Supplemental Table 6). This confirms that the lentiviral integrations in the immortalized cell lines did not cause an overt genomic difference compared the matched primary culture genomes.

**Figure 3.**
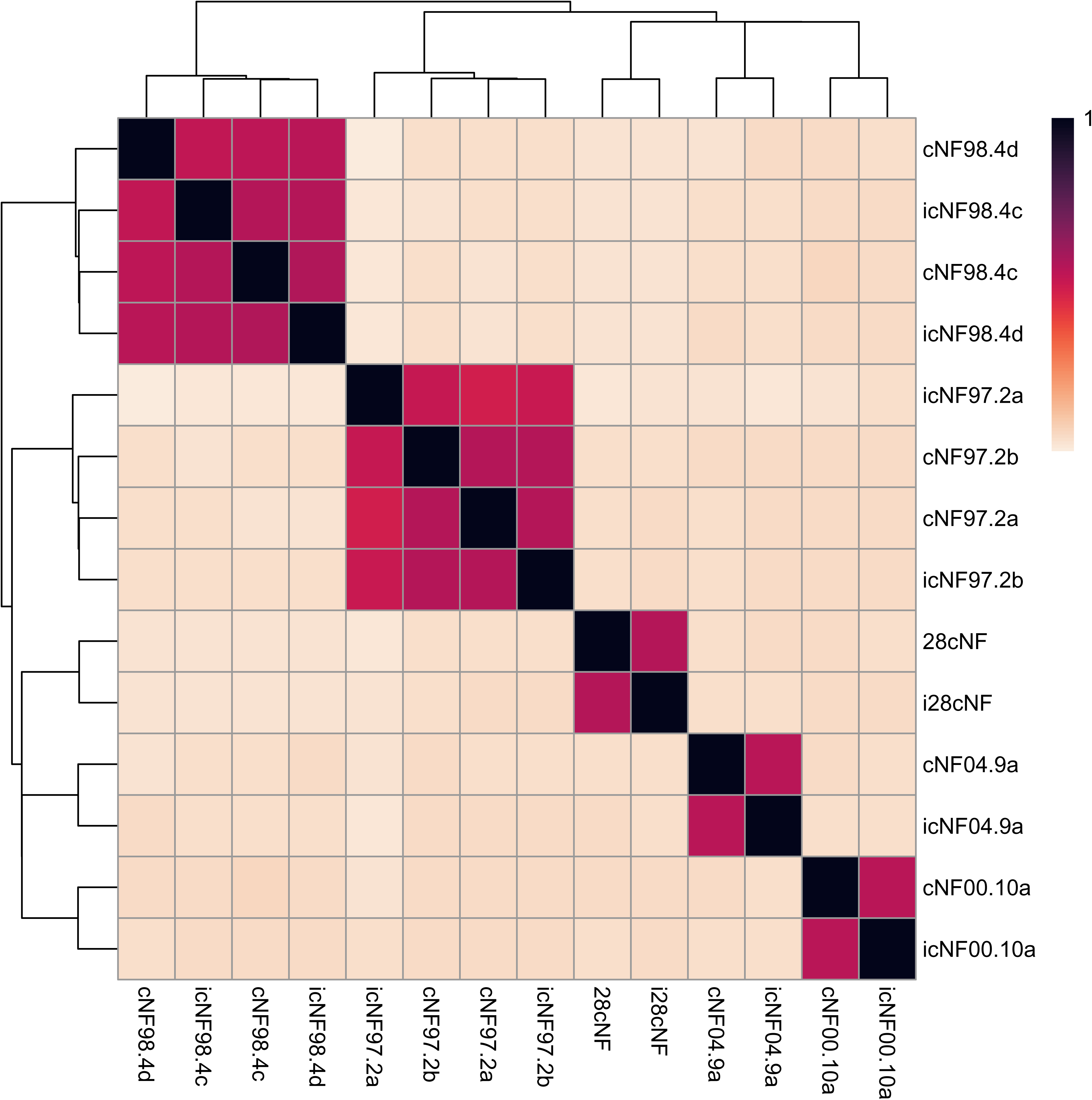
Hierarchical clustering of Jaccard indices. The Jaccard index, a measure of similarity, was calculated for all cell lines in a pairwise manner using the variants called with DeepVariant. A value of 0 indicates no intersection between the called genomic intervals (and thus no similarity), while 1 indicates perfect concordance. The cell lines that are immortalized/primary pairs (Jaccard index ranging from 0.679 to 0.722) cluster together (red and black boxes), indicating that the immortalized cell lines have a variant profile that most closely matches that of the corresponding primary cell culture.

CNAs were assessed in the WGS data and compared to gene expression quantification (Supplemental Figures 3-16). Regions of missing data or small amplifications are also observed, however, as seen in Supplemental Figures 3-16 these typically correspond to centromeric regions. We did not observe evidence of CNAs that occurred across all cNF cell lines, or evidence of copy number alterations associated with hTERT/mCdk4-mediated immortalization.

### Transcriptomic profiling of primary cell cultures and immortalized cell lines

Principal component analysis of transcriptomic data showed weak grouping of the primary cell lines and matched immortalized cell lines (Figure 4A). Principal components 1 and 2, which captured the largest sample variance based on expression profiles, only accounted for 21.6% and 18.7% of the variance between samples, and both of these components correlated with immortalization status. The Euclidean distances between cell lines/cultures were calculated from complete gene expression profiles, and the pairwise distances were hierarchically clustered (Supplemental Figure 17). Most immortalized cell lines clustered closer to one another than their paired primary cell line (icNF00.10a being an exception). However, when compared to *NF1* plexiform neurofibroma cell lines immortalized using the same methodology,^20^ the gene expression distances between primary cultures and immortalized cell lines were much smaller in magnitude than observed between the tumor types.

**Figure 4.**
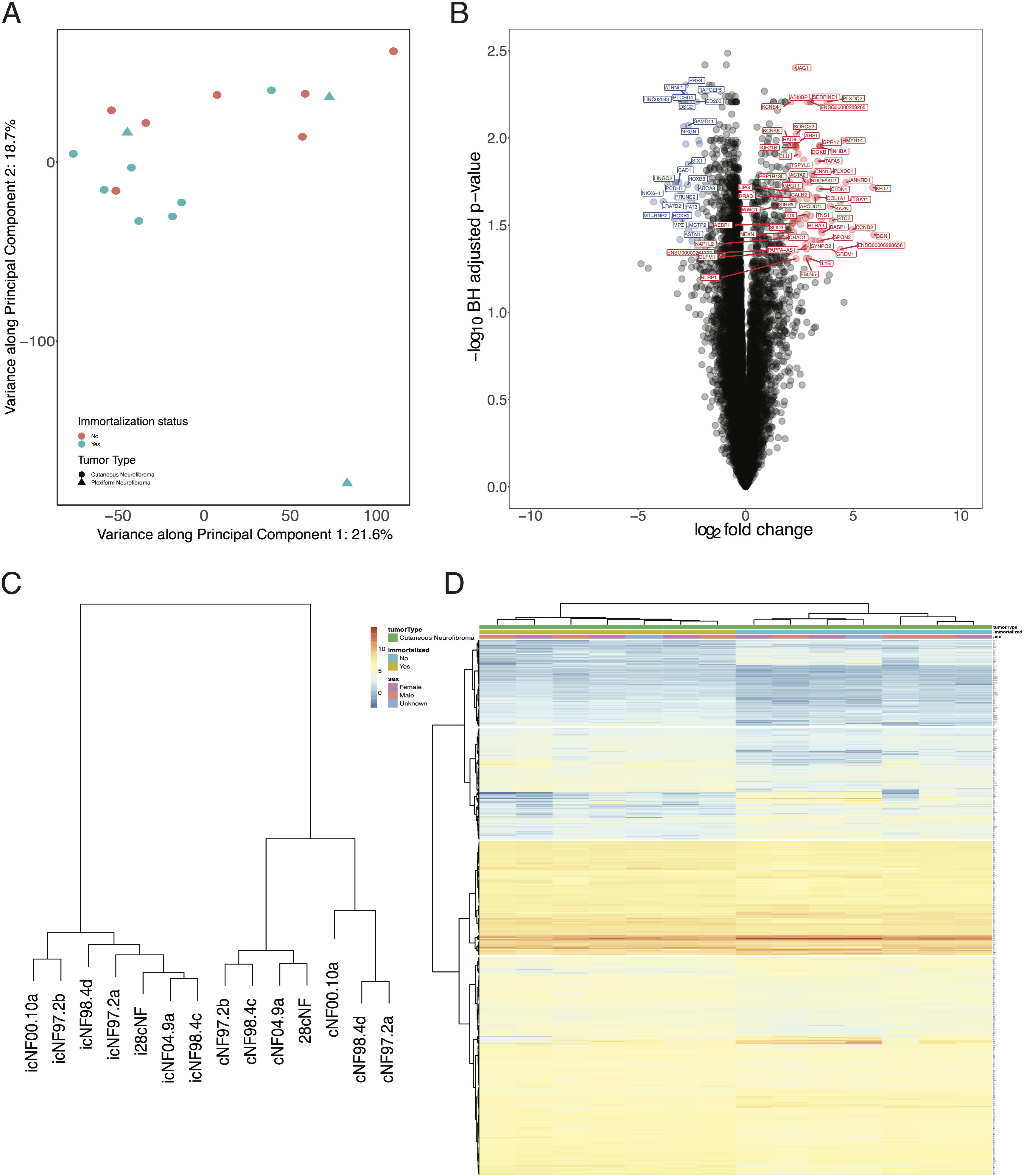
Transcriptome profiling results of immortalized cell lines compared to primary culture counterparts. (A) Principal component analysis of cell lines indicates minimal grouping by either immortalization status or tumor type; representative pNF cell lines show greater variance across PC2 than the cNF cell lines. (B) Volcano plot showing overview of differentially expressed genes in immortalized versus primary cells. Blue points indicate genes that have significantly increased expression in immortalized cNF cells (log2 fold change > -2, BH-adjusted p-value <0.05). Red points indicate genes that have significantly increased expression in primary cNF cell cultures (log2 fold change > 2, BH-adjusted p-value <0.05). (C) Hierarchical clustering (WardD2 clustering) showing sub groups of primary cell cultures and immortalized cell lines based on. (D) Heatmap showing expression (logCPM) of differentially expressed genes between immortalized and primary tumor cells.

Gene expression profile analysis found significantly different transcript levels between primary cultures and immortalized cells, for 993 genes (Figures 4B and 4C), with four sub-clusters based on differentially expressed genes (Figure 4D). Pathway enrichment analysis of the differentially expressed genes suggested the perturbation of cell cycle regulatory pathways (e.g. ‘regulation of cyclin-dependent protein kinase activity’, ‘regulation of G1/S transition of mitotic cell cycle’), consistent with mCdk4 and hTERT-mediated cellular reprogramming to the immortalized state (Supplemental Table 6). *NF1* gene expression did not significantly differ between matched primary and immortalized cell lines, indicating preservation of *NF1* status (NF1: log fold change = -0.02, adjusted p-value = 0.96).

## Discussion

cNF are the most common tumor in people with NF1. They cause significant morbidity and disfigurement impacting quality of life for the vast majority of people living with NF1. Although they share some similarities with other neurofibromas, they have distinct behavior and in the absence of any approved systemic or topical therapy available, are in great need of therapeutic discovery efforts. Enriched SC cultures from primary cNFs are not often completely pure; 1-10% of the cells are of other types, most commonly fibroblasts. This, and the limited passages prior to senescence, are barriers to many potential preclinical studies. We previously had success creating immortalized SC lines from NF1 pNFs and wild-type human SC, overcoming those barriers, with the caveat that the cells were genomically modified.^20^ We determined that transduction with both hTERT and mCdk4 cDNAs was needed to enable the pNF SC to pass beyond p10-20. These cell lines have since been used in studies such as drug screening and xenograft experiments.^63–65^ Given this success, we used the same approach to immortalize cNF primary SCs with the goal of accelerating drug discovery efforts for this tumor. Out of the 15 attempted co-transductions, 14 were successful. These were “population” co-transductions of 10,000 cells, with no specific selection. Although the primary cultures were developed with conditions to enrich for two-hit SCs, many still have a minor component of heterozygous SC and stromal cells such as fibroblasts. The resulting cell lines (considered successful if divided for more than 6 passages beyond the primary culture lifespan) thus could contain immortalized cells other than two-hit SC. In future studies, approaches such as a panning enrichment for p75 positive SCs may be a possibility to remove non-desired fibroblasts. Additional variability could come from the number and location of transgene integrations, and any subsequent genome alterations.

Unlike the immortalized pNF cell lines, some of the icNF lines either senesced or slowed in proliferation rate before passage 40, and nearly all require or prefer a substrate and neuregulin growth factor, suggesting that they are only semi-immortalized. This could be the result of intrinsic differences between cNF and pNF SC, or poorer transduction (done in 2 batches). Consistent with this, of the 12 icNF cell lines for which we obtained cytogenetic data, the karyotypes were all normal (46,XX or 46,XY) except that icNF18.1a (p21) also had a somatic balanced reciprocal translocation (t(15;22)(q22;q13)), and 10% of icNF97.2b (p14) cells were tetraploid with no rearrangements. Tetraploidy was seen among a number of the immortalized plexiform SC lines created previously,^20^ which could be due to high expression of mCdk4.^66^ Presumably, in the early passages after transduction, a number of cells acquired and expressed the transgenes, with the most robust clones continuing throughout culturing. We can’t determine how many clones grew out to establish the cell line, but it is likely at least several (because not all of the icNF cells had the same characterizations), and these clones may still be competing in culture. Only icNF97.2b was mosaic at the cytogenetic level. The presence of multiple clones can actually be an advantage, providing an opportunity to isolate different lines via single-cell cloning (something that cannot be done with primary SC). We did this to obtain a matched heterozygous icNF line from icNF98.4, and previously used single-cell cloning to isolate a two-hit SC line from an immortalized plexiform cell line that appeared mostly heterozygous at *NF1* (ipNF08.1).

The presence of a variety/combination of *NF1* germline and somatic pathogenic variants in these cell lines may also be advantageous for groups investigating the effects of specific variant types or locations on function. While this genetic heterogeneity, and the heterogeneity in other cell measures, adds complexity to future studies with these cell lines, this cohort is consistent with the significant molecular and clinical heterogeneity in human NF1 tumors.

Interestingly, there was no statistical difference between pERK:ERK ratios of cell lines that lacked full-length neurofibromin versus those that expressed at least some apparently-full-length neurofibromin. A consideration is the possibility of function from truncated neurofibromin molecules, a phenomenon not addressed here, nor yet proven in NF1. However, analysis of neurofibromin isoforms containing truncating nonsense variants in a cDNA-based overexpression system suggest that isoforms truncated after the GRD may still retain the ability to suppress Ras activity.^67^

Bioinformatic analysis of WES data indicated that the icNF cells were genetically closer to their matched primary cells than cells from other patients, and that the majority of expression differences are related to the cell cycle, consistent with transgene function to extend lifespan. One cell line (icNF97.2a) has already been published in a study by our group, chosen for its specific mutation repertoire to test an exon skipping approach.^68^ This cell line as well as 2 others (icNF97.2b and icNF98.4d) have also been published by our group to evaluate the transcriptomic effects of selumetinib.^69^ Interpretations about some cell lines are not firm, such as whether one germline variant is pathogenic, and regarding the cell lines that appear genetically heterozygous (Table 1). Additional cellular and biochemical characterizations were beyond the scope of the grant supporting this work, but will hopefully be taken up by others in the future.

This work describes the first immortalized SC lines from human NF1 associated cNF. The majority of the cell lines contain the *NF1* two-hit tumor cells. These, plus the heterozygous cells, will contribute to research in tumorigenic mechanisms and therapeutic development for these common and yet largely unaddressed tumors.

## Acknowledgements.

We thank the patients and their healthcare providers who contributed the original tissue samples and clinical data. This publication was supported by a subagreement from the Johns Hopkins University via the Neurofibromatosis Therapeutic Acceleration Program (NTAP) with funds to Dr. Wallace provided by a Grant Agreement from the Bloomberg Family Foundation. This publication’s contents are solely the responsibility of the authors and do not necessarily represent the official views of the Bloomberg Family Foundation or the Johns Hopkins University. Other support for this work includes: Gilbert Family Foundation’s Gene Therapy Initiative (Dr. Wallis); the Intramural Research Program of the Division of Cancer Epidemiology and Genetics, National Cancer Institute, and National Institutes of Health (Drs. Stewart and Pemov); NTAP cNF initiative project (Dr. Serra); NTAP Data and Knowledge Portal for NF1 (Dr. Allaway and Dr. Banerjee). The authors thank Ms. Kristin Jones (Frederick National Laboratory for Cancer Research, Division of Cancer Epidemiology and Genetics, National Cancer Institute) for supervision of *NF1* high-throughput sequencing and Dr. Jung Kim for assistance with AutoGVP.

## Author Contributions

H.L, L-J.C., A.P., D.M., J.L., J.N., M.C., H.M., X.Z., J.B., R.A., A.S., D.W. and Y.L. generated materials or data for this publication experimentally, and/or contributed bioinformatic and statistical analysis. M.W., E.S., J.O.B., A.H., A.Y., D.S. and D.W. contributed to the concept, design of the work and/or data interpretation. M.W., H.L., A.P., J.B. R.A., D.S., D.W., and A.H. participated in the writing of the manuscript. All authors read and approved the final paper, with the opportunity to contribute revisions.

## Abbreviations

mCdk4: murine cyclin dependent kinase 4
NF1: neurofibromatosis type 1
SC: Schwann cell(s)
hTERT: human telomerase reverse transcriptase
cNF: cutaneous neurofibroma

