## Supplemental Figures for "Immortalization and Characterization of Schwann Cell Lines Derived from NF1 Associated Cutaneous Neurofibromas"

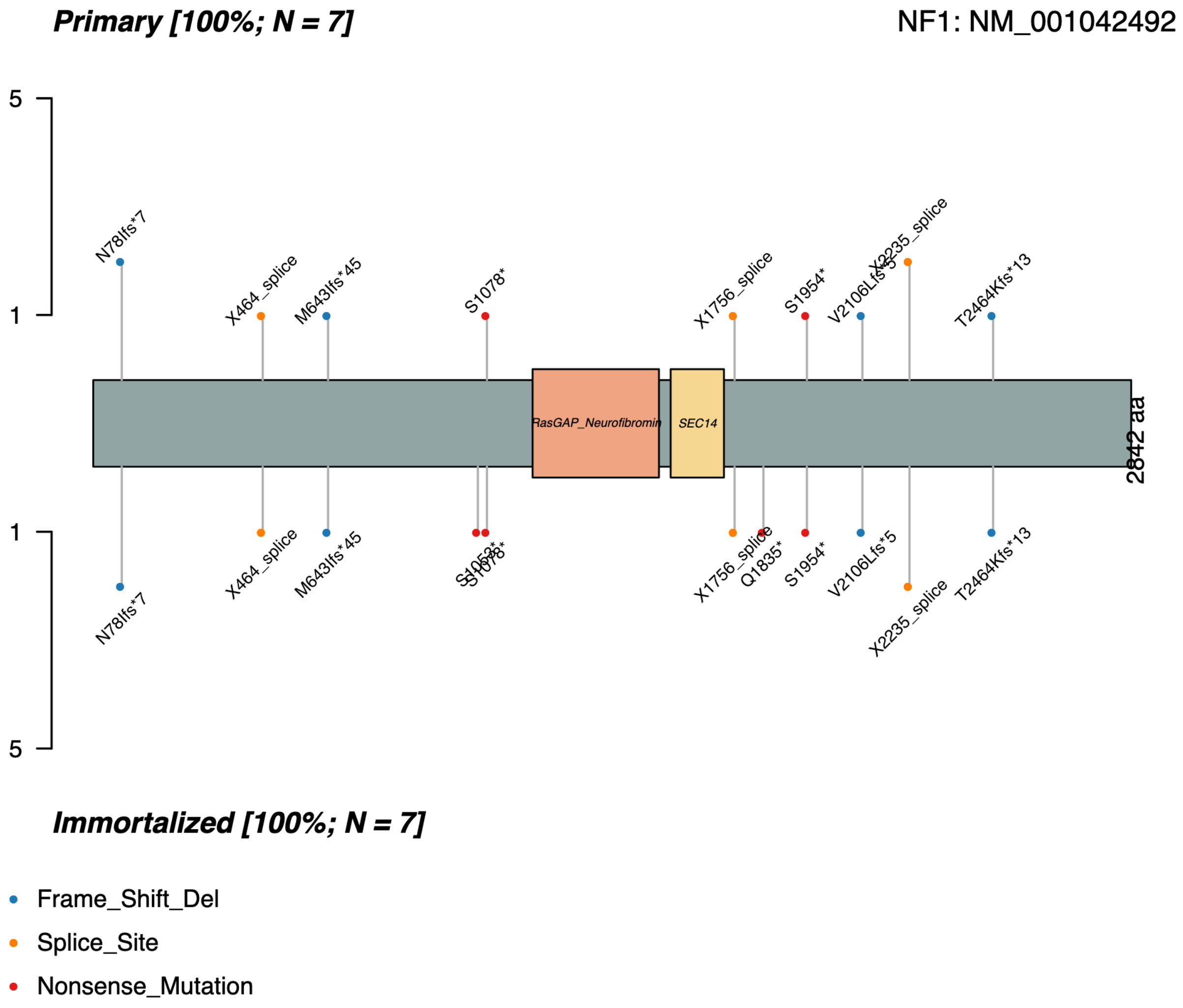


Supplemental Figure 1 - **NF1 mutations detected in the primary (top) and immortalized (bottom) cNF cell lines from whole-genome sequencing data.**

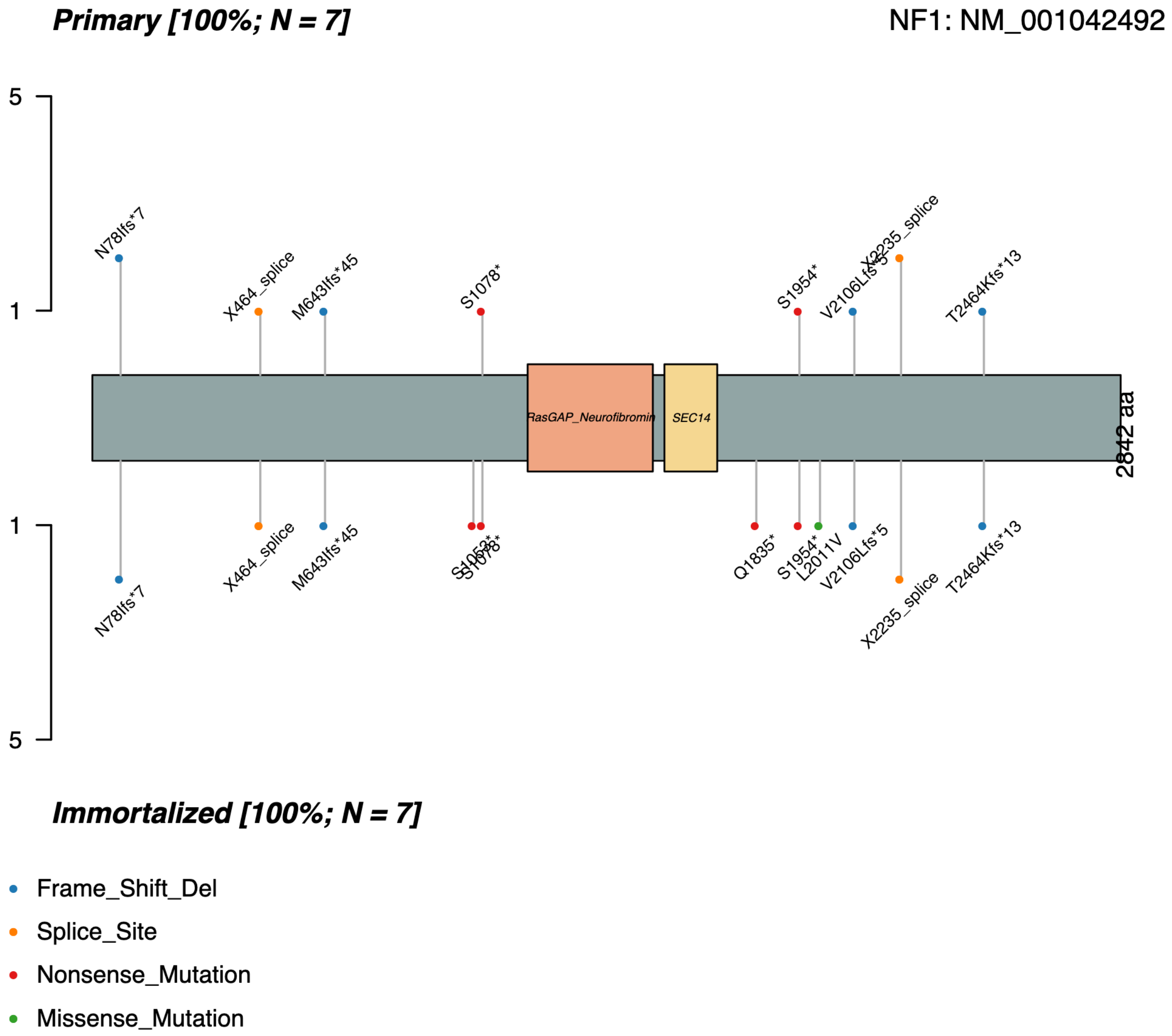


Supplemental Figure 2 - **NF1 mutations detected in the primary (top) and immortalized (bottom) cNF cell lines from whole-exome sequencing data.**

Supplemental Figures 3-16 (starts next page): **Copy number variation and associated gene expression in all cell lines profiled by whole genome sequencing.** Circos plots show the copy number variation across the profiled genome (red) and the log 10 transformed gene expression at each of those locations. As expected, telomeric and centromeric regions have reduced transcription/associated gene expression, as do Y chromosome genes in cell lines that have no copies of this chromosome. However, other large "deletions", such as chr4 in cell line 28cNF do not lack transcription of genes in these regions, suggesting that these deletions are likely an artifact. This analysis demonstrates that the sex of each cell line was as expected, and that the immortalization process did result in large copy number alterations in the cNF cell lines.

### 28cNF

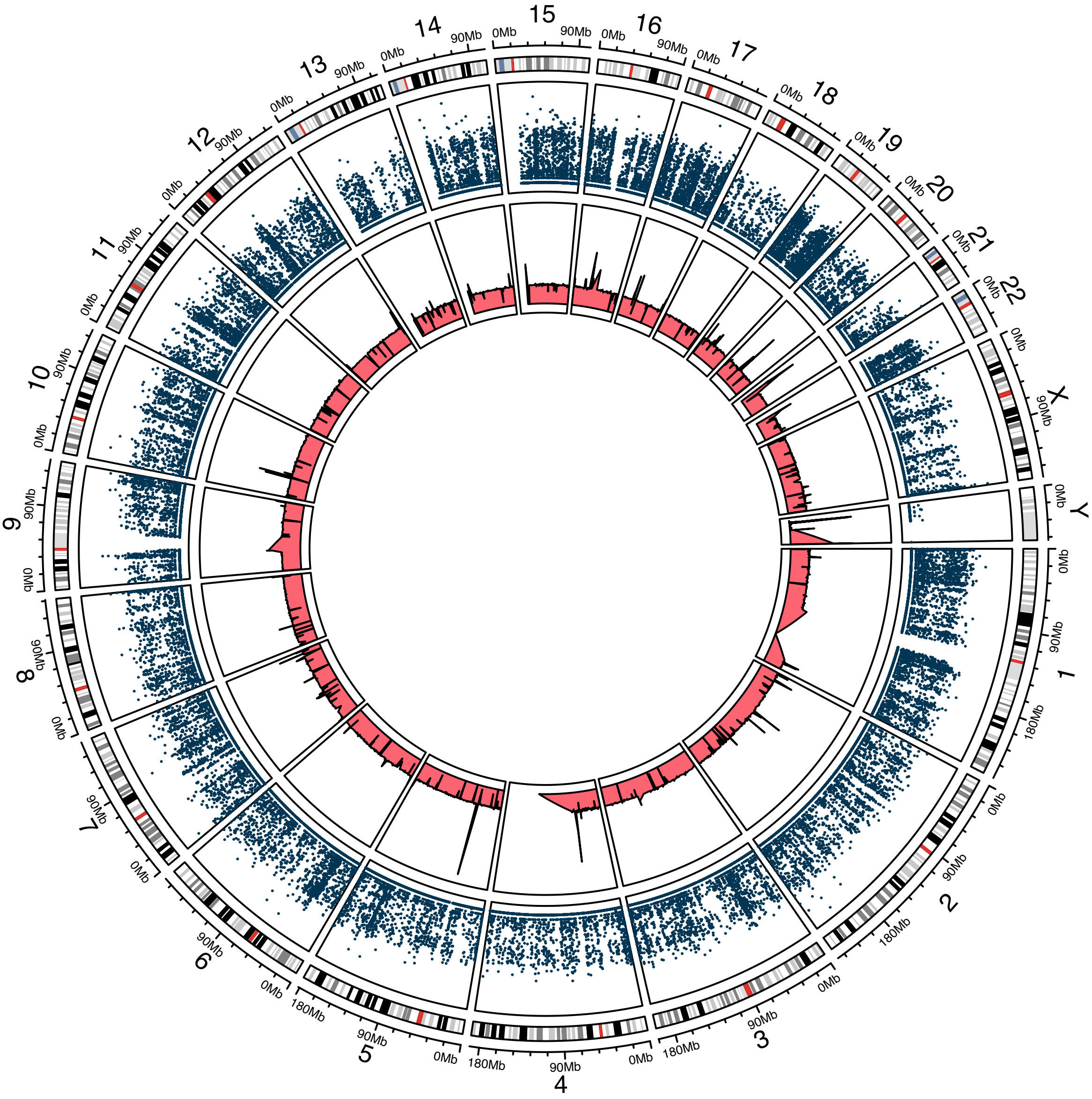

### i28cNF

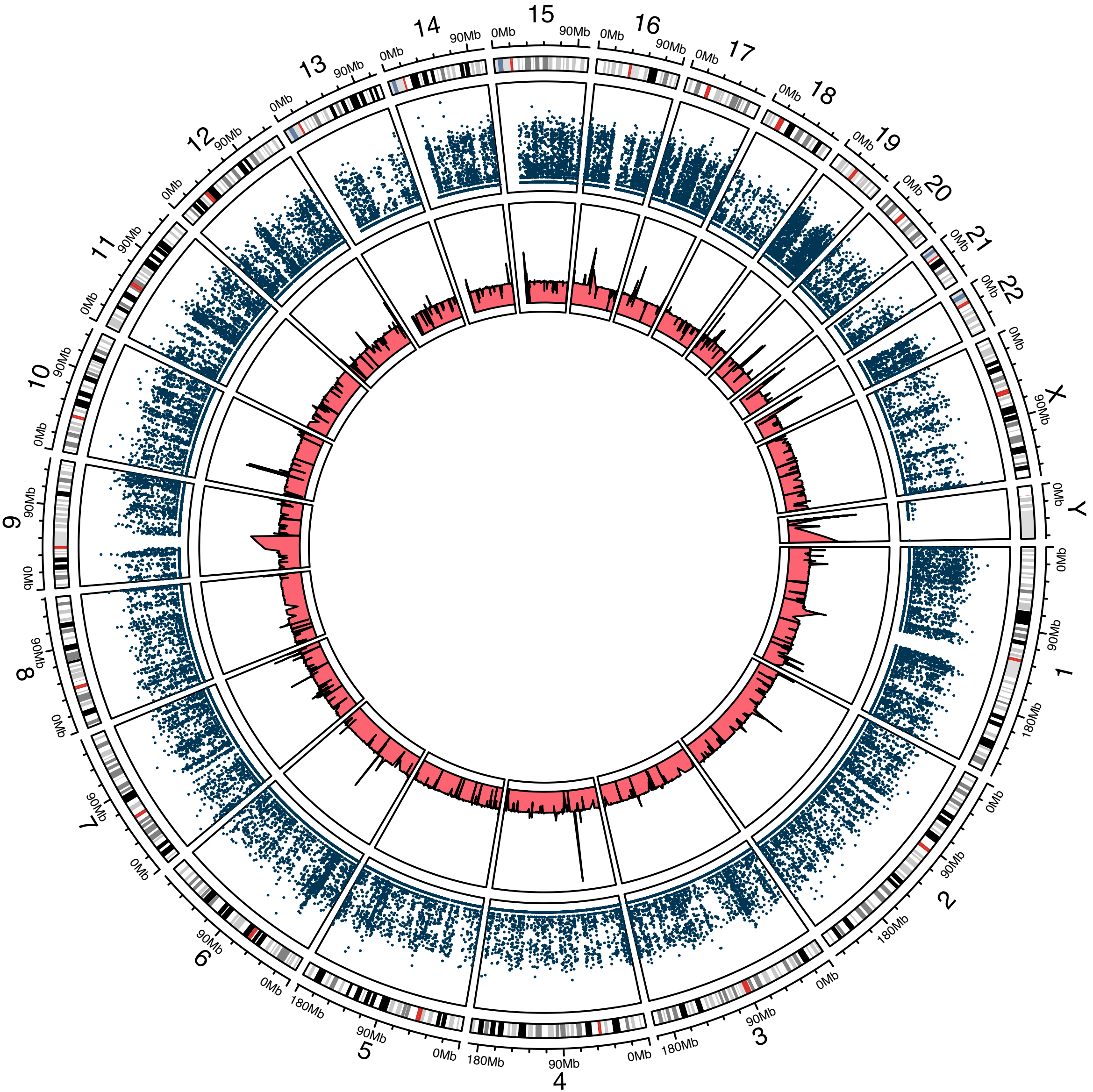

### cNF00.10a

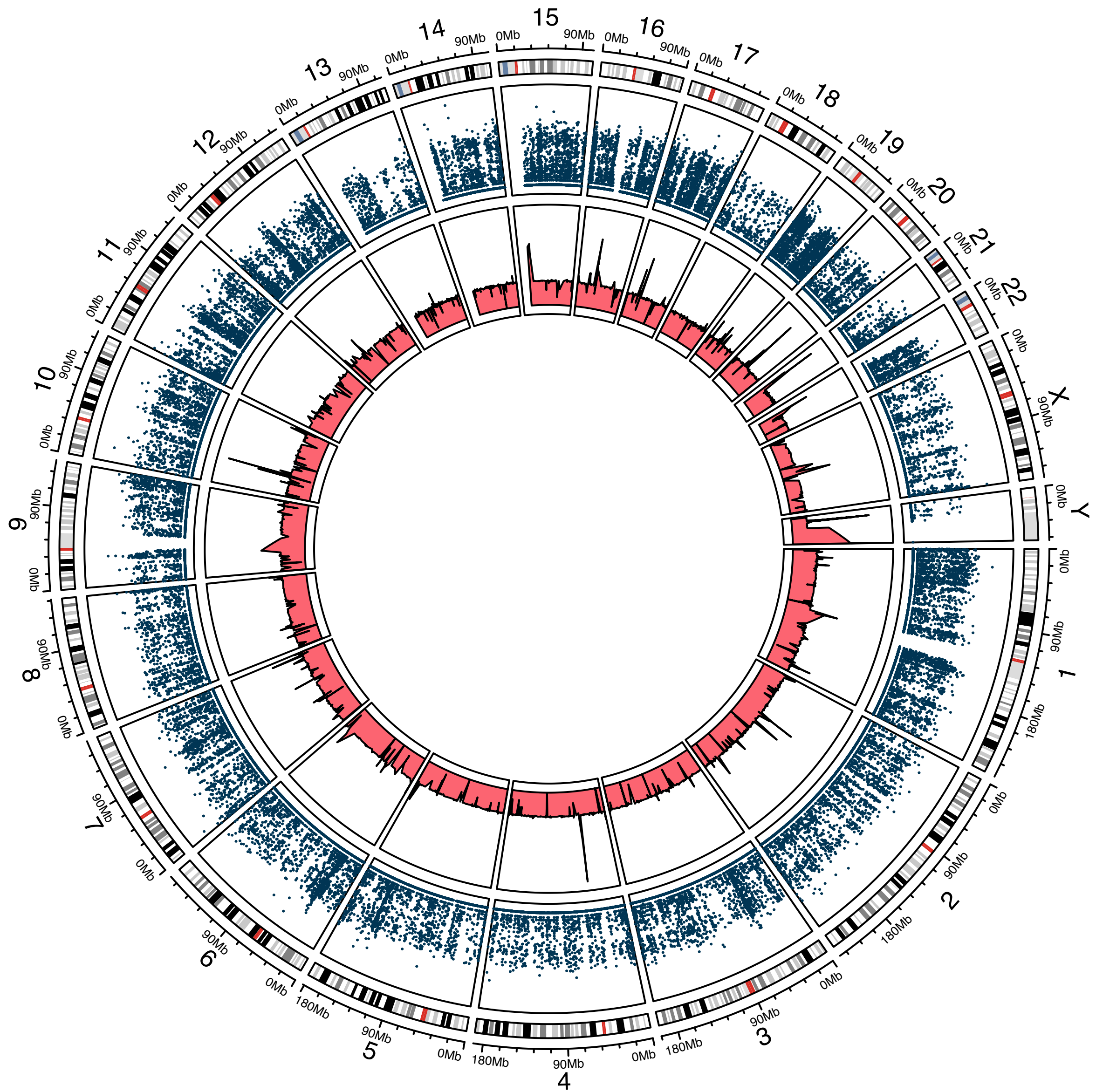

#### icNF00.10a

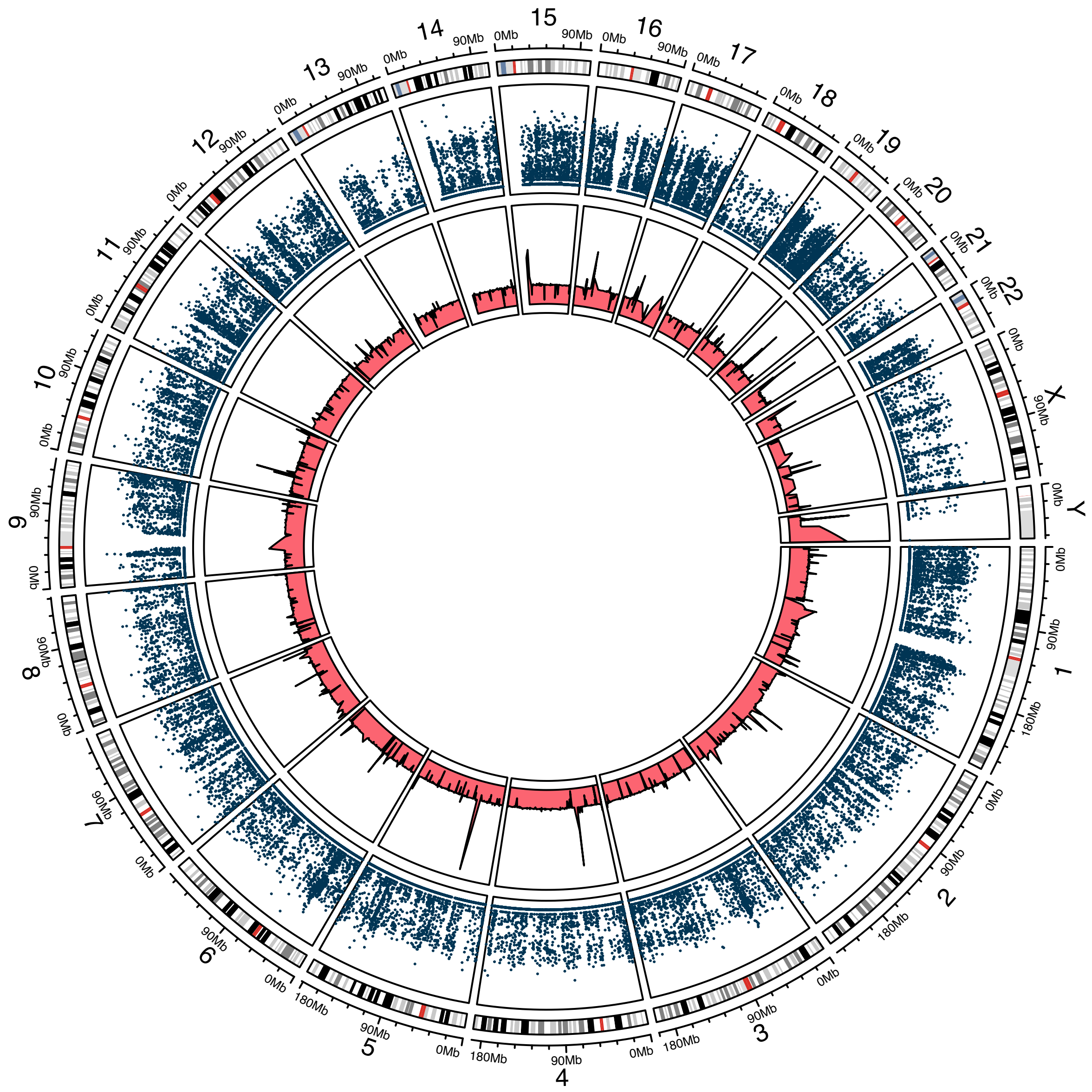

### cNF04.9a

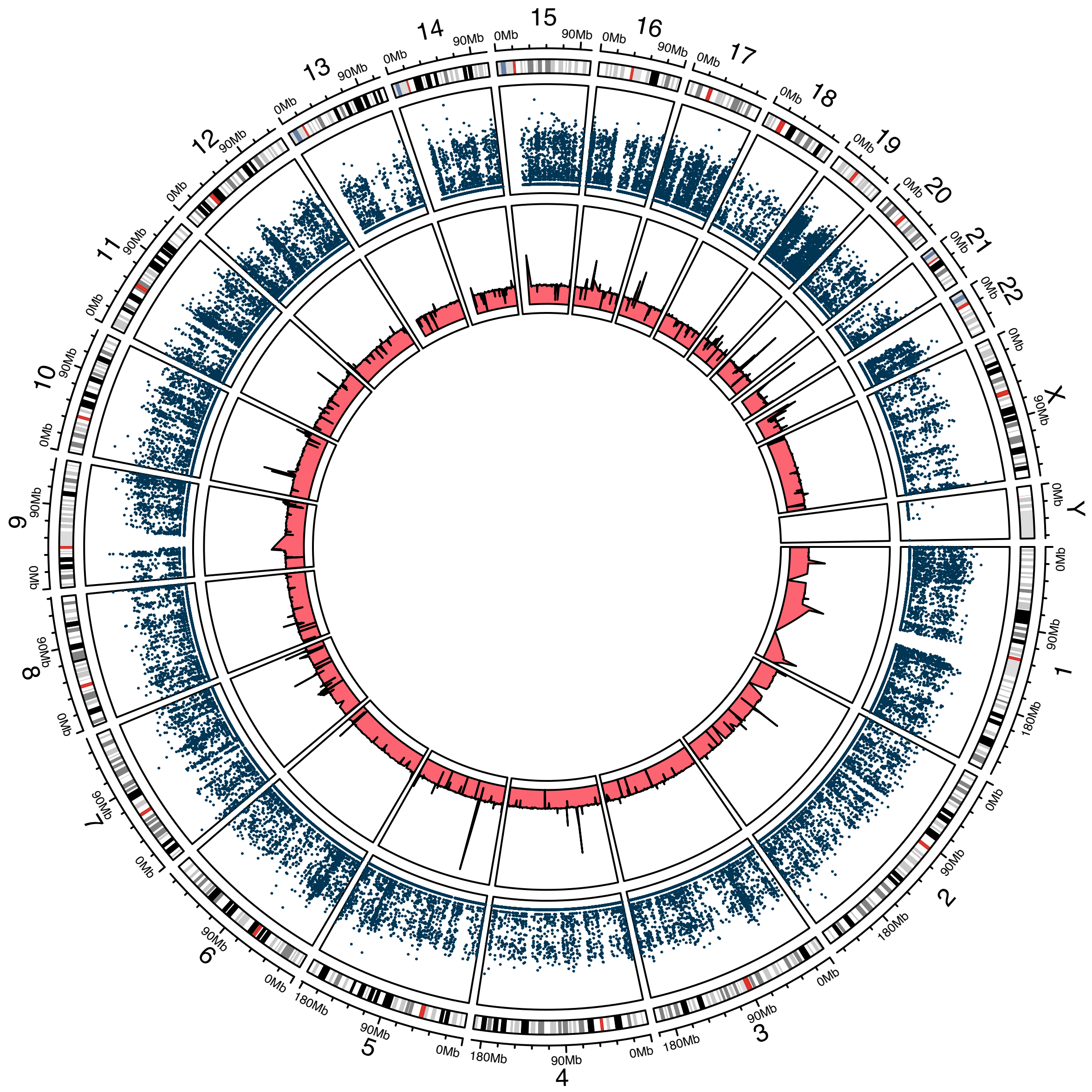

### icNF04.9a

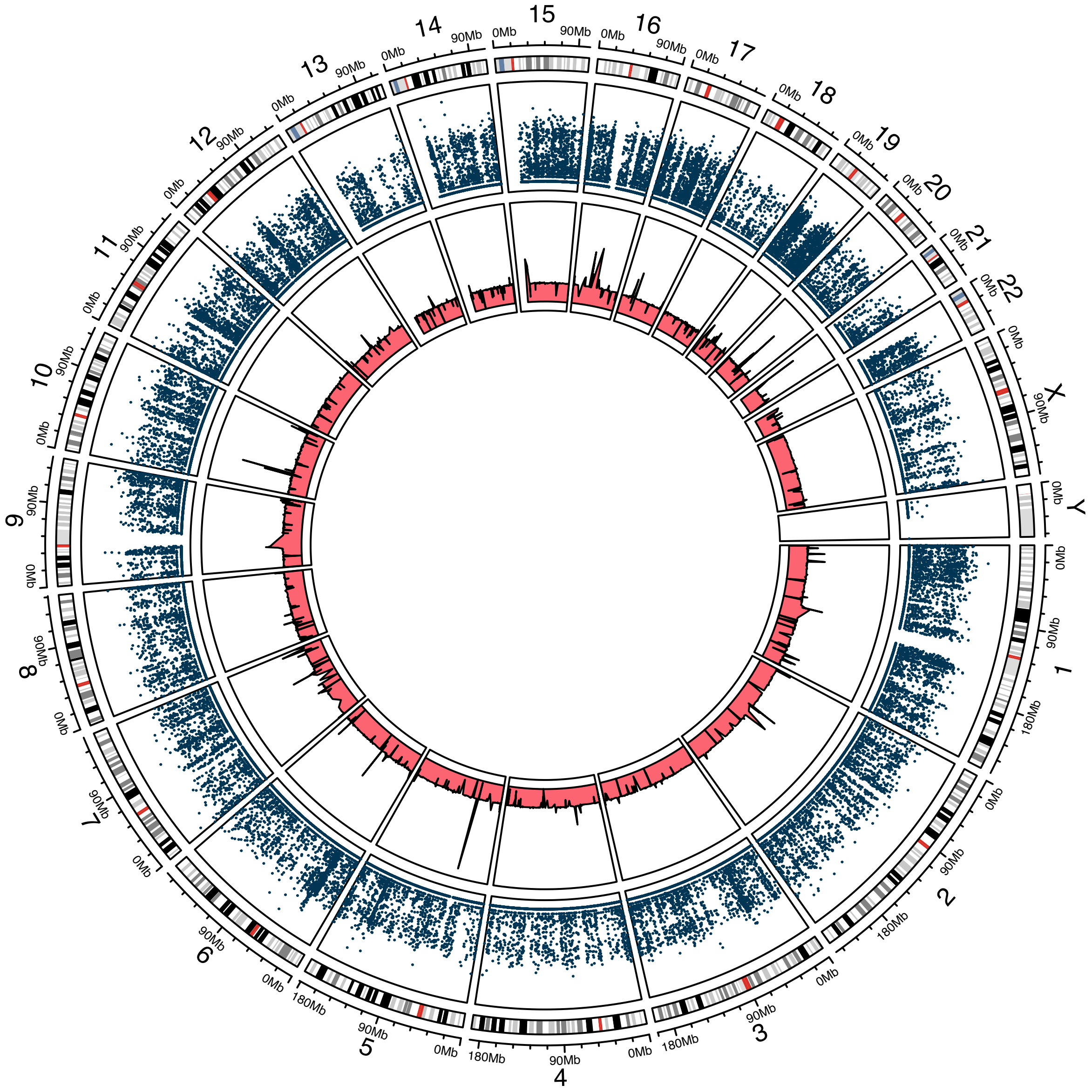

### cNF97.2a

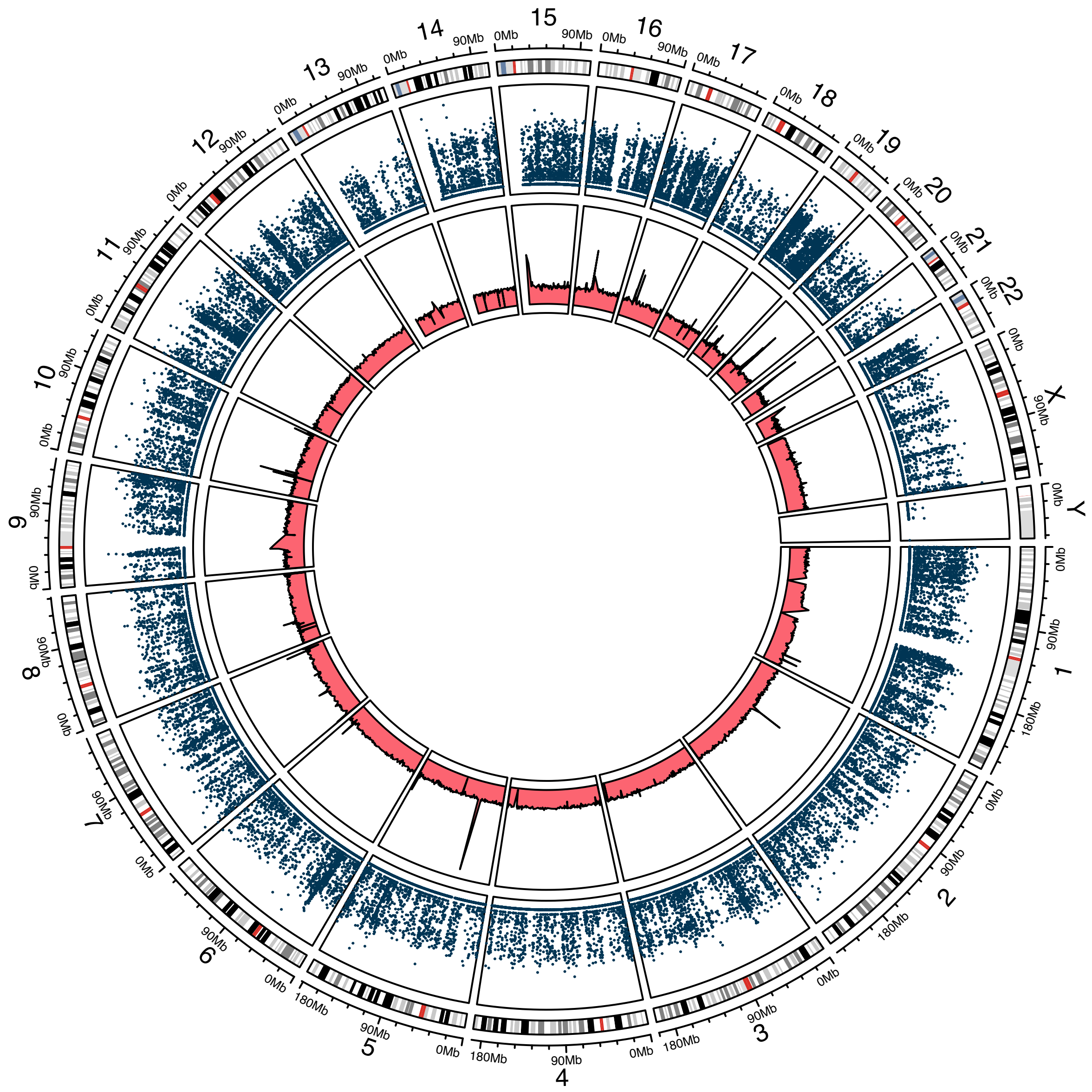

### icNF97.2a

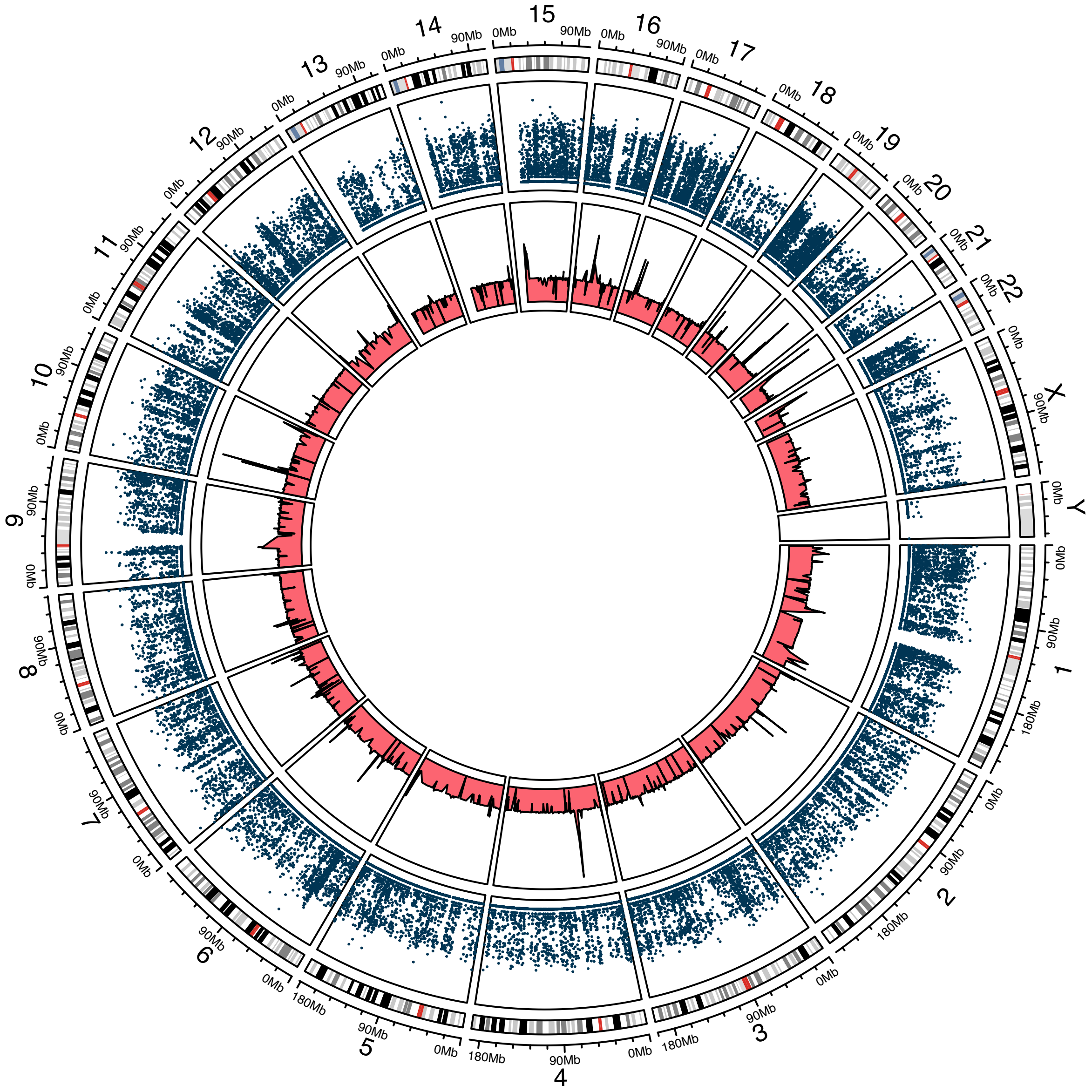

### cNF97.2b

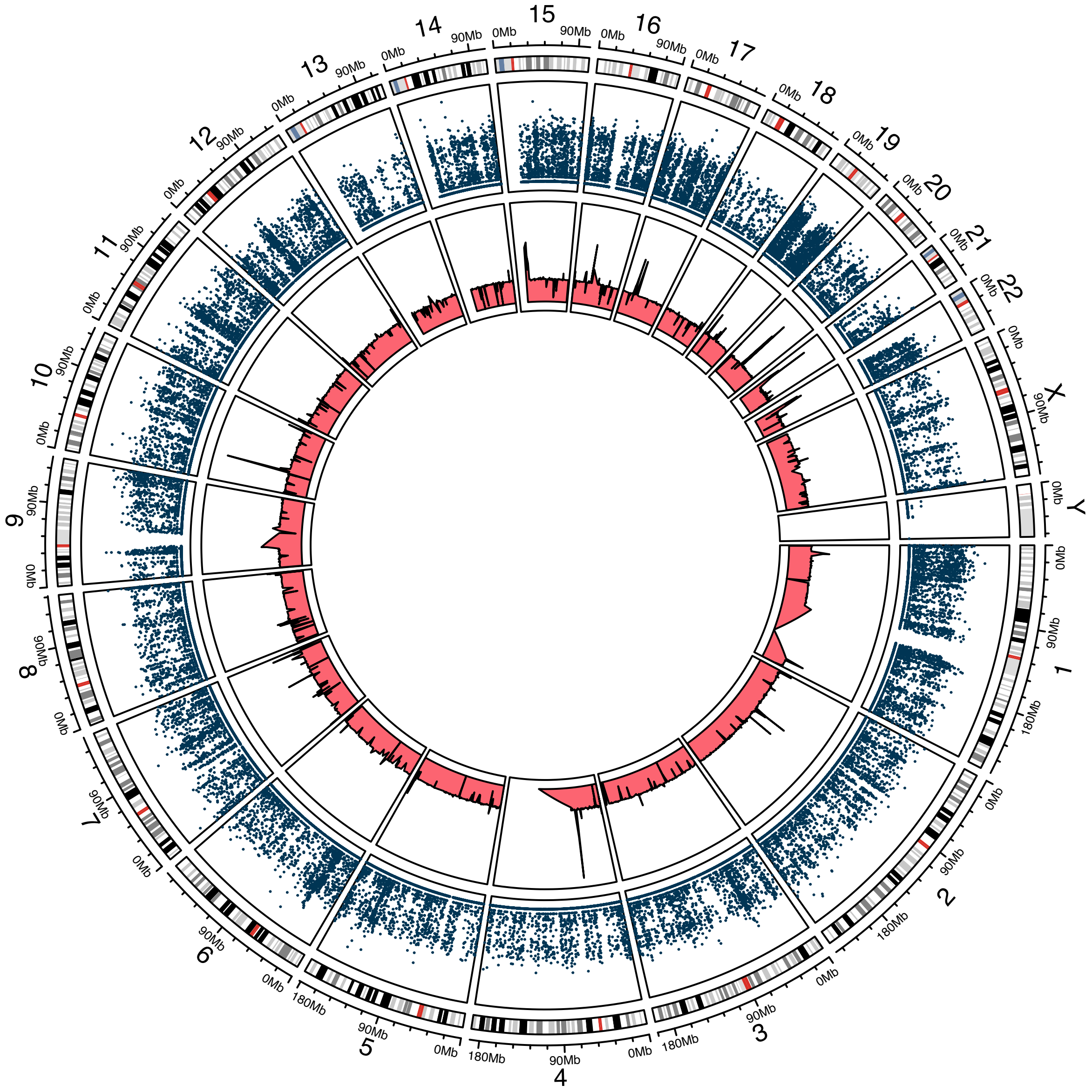

icNF97.2b

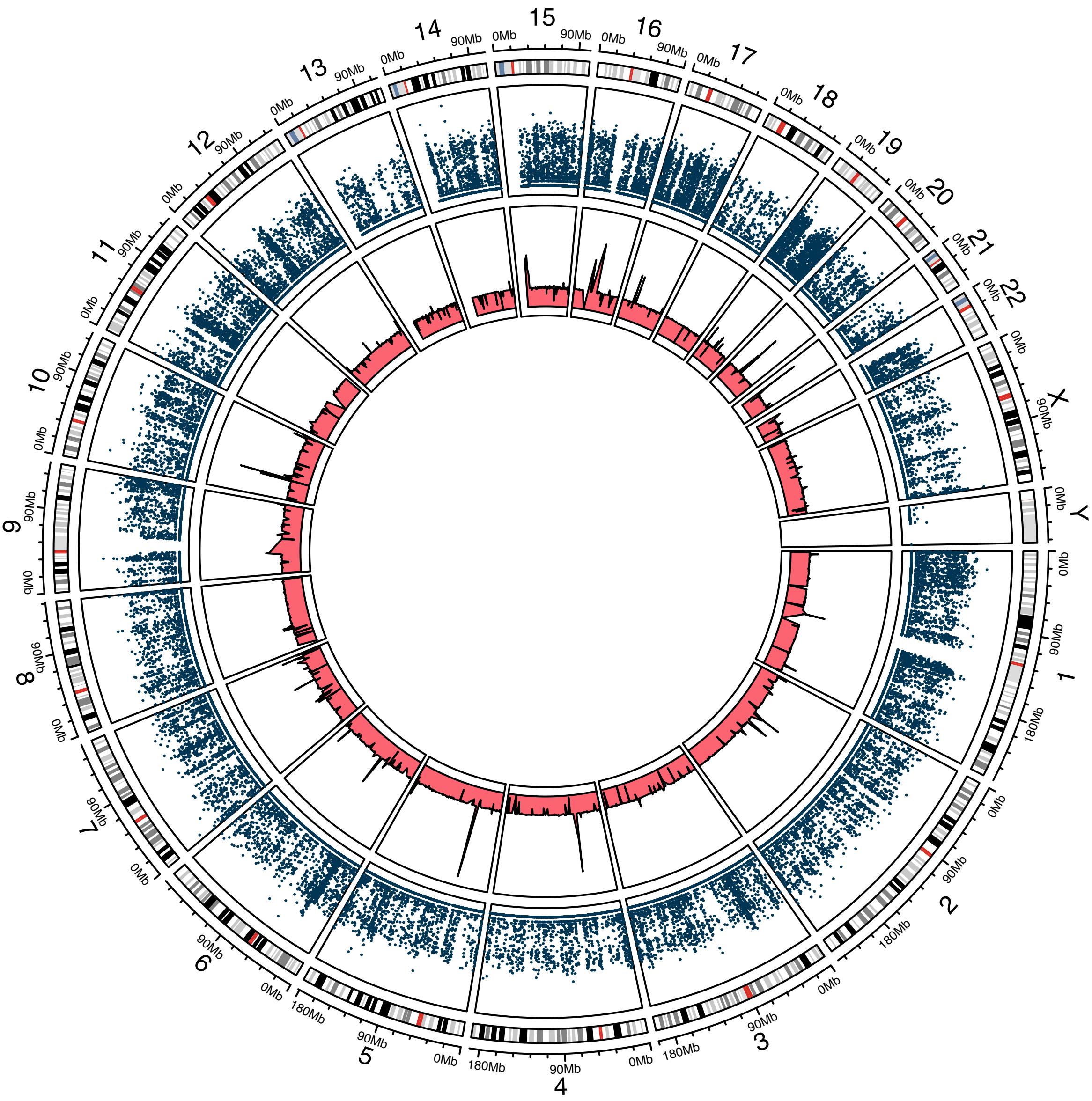

### cNF98.4c

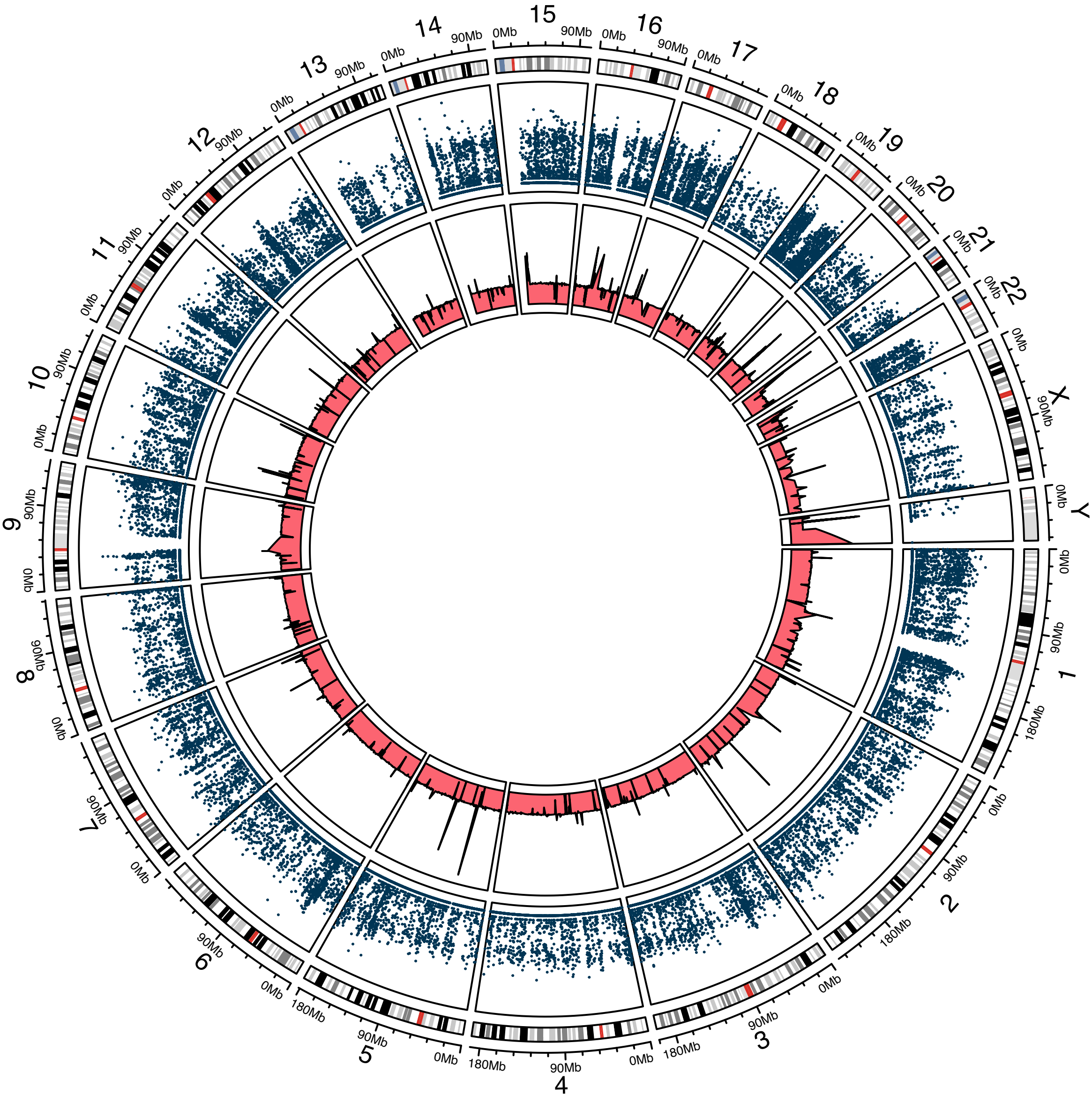

### icNF98.4c

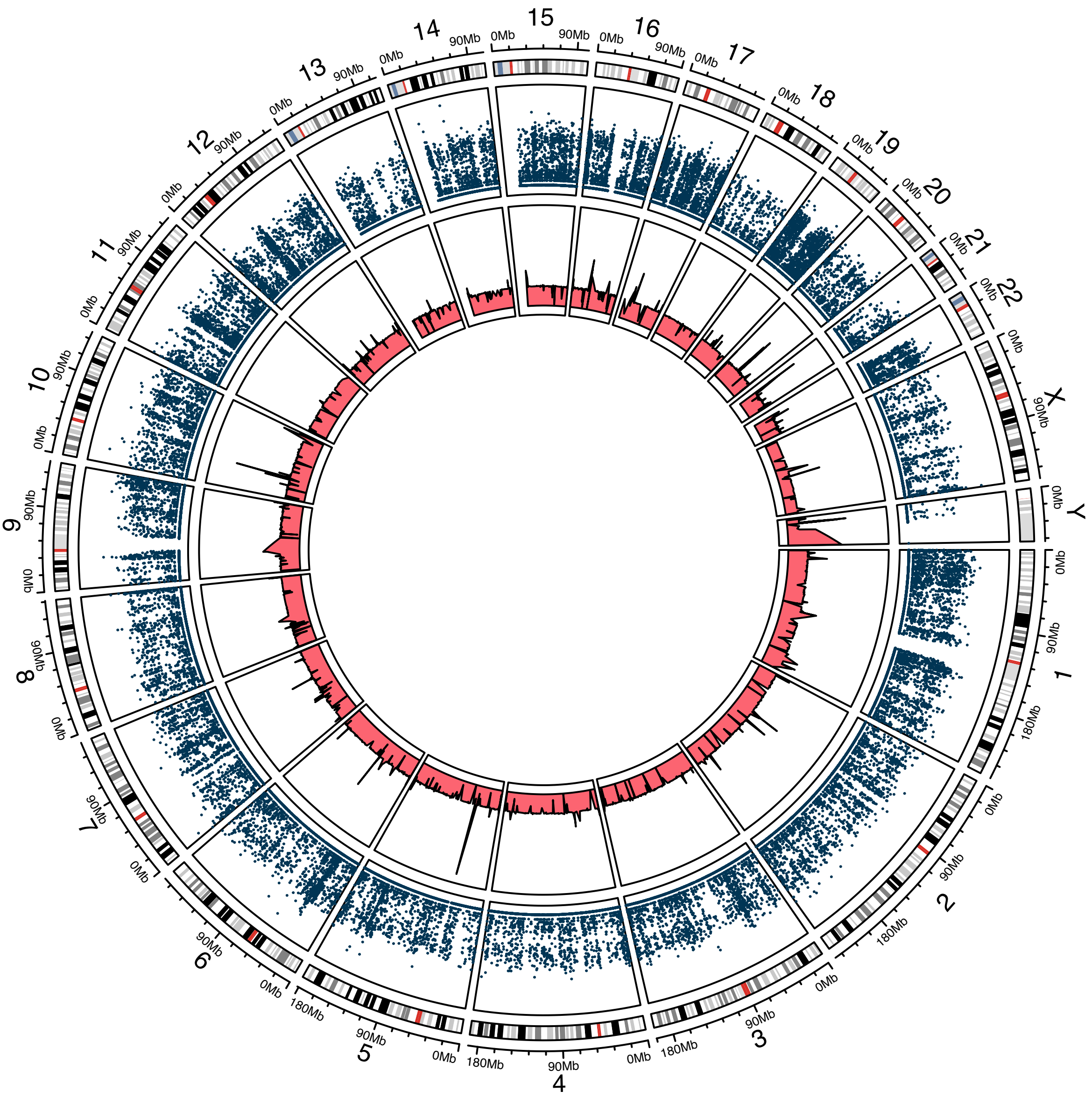

#### cNF98.4d

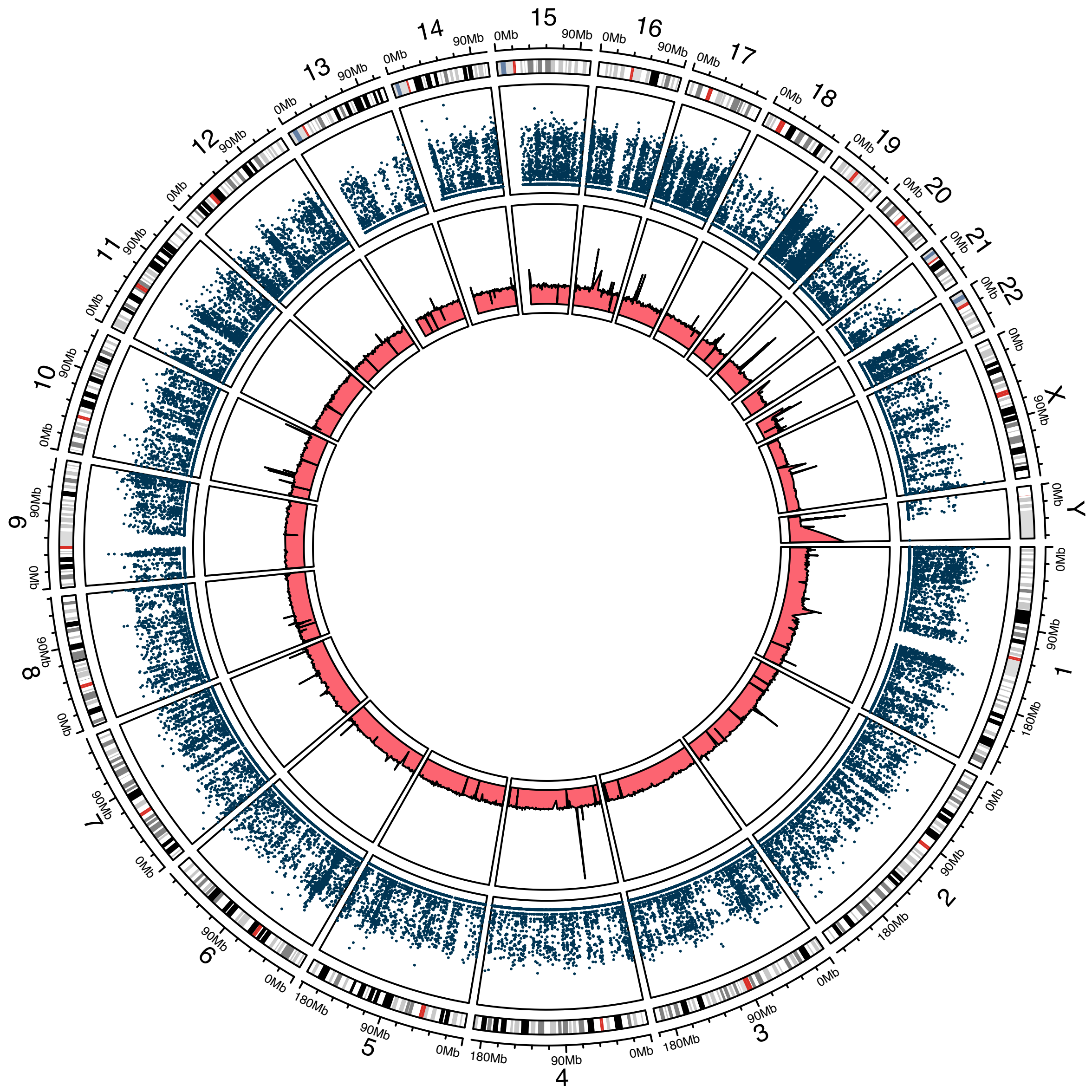

icNF98.4d

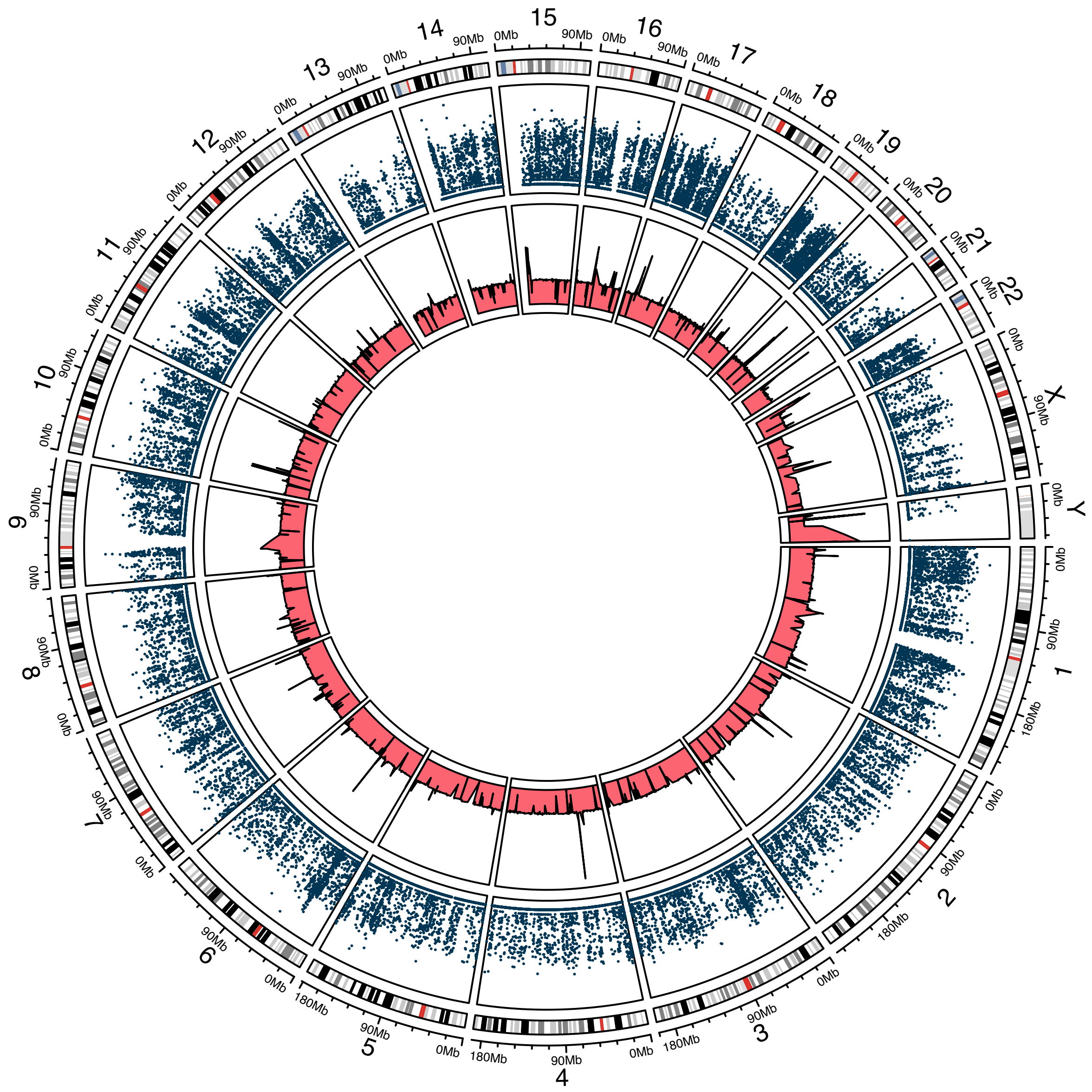

Supplemental Figure 17. **Euclidean distances between primary cultures and immortalized cell lines based on complete gene expression profiles.** Hierarchical clustering of the Euclidean distances show that the transcriptome profiles of primary cultures from the same patient and their immortalized cell lines were more similar (lower distance) to each other than to that of primary cultures and immortalized cell lines from other patients or other tumor types (e.g. plexiform neurofibromas: hTERT_NF1_ipNF95.11b_C and hTERT_NF1_ipNF95.6).

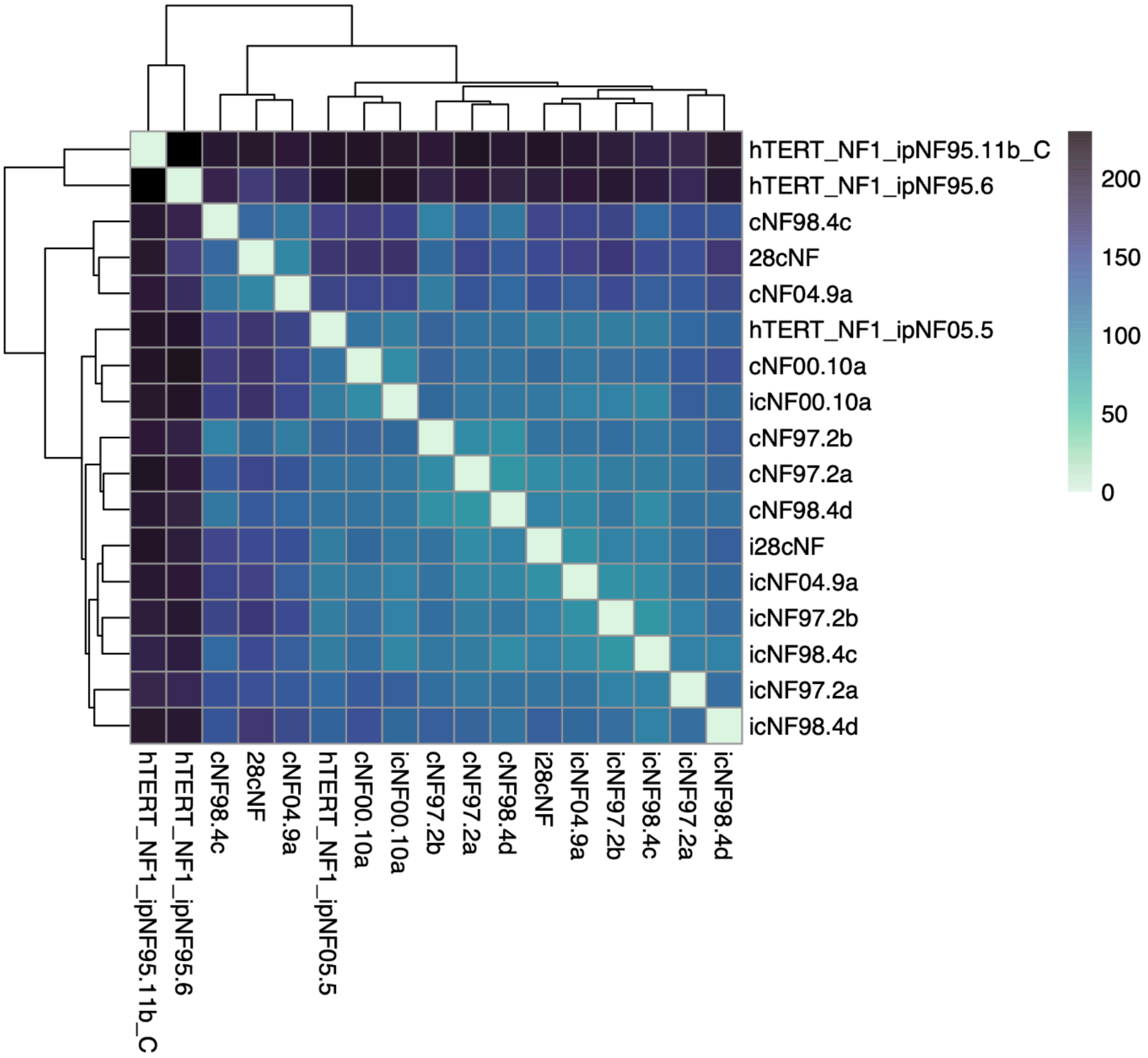
