## Supplemental Table 2 for "Immortalization and Characterization of Schwann Cell Lines Derived from NF1 Associated Cutaneous Neurofibromas"

Supplemental Table 1. NF1 RT-PCR primers for Sanger sequencing

| Primer name | Sequence (5’ > 3’) | Exon* | Amplicon size (bp) |
| --- | --- | --- | --- |
| NFC.1F | CTCTCCTTGCCTCTTCCCTCA | 1 | 393 |
| NFC.1R | CCCAGCAAGACATTTTTCCAGTG | 3 |  |
| NFC.2F | TGGTTATAAGCGGCCTCACTA | 2 | 533 |
| NFC.2R | TTCATCTGGATAATTTTCTACCCAGTTC | 7 |  |
| NFC.3F | CGACTCCTGAAGGAAACAGCA | 6 | 526 |
| NFC.3R | CTCTTGAGAATGGCTTACTTGGATTAAA | 10 |  |
| NFC.4F | CTGGCCATGGAGGAAGTAGG | 9 | 458 |
| NFC.4R | TATTGCTGGGTGTGCTCCAC | 12 |  |
| NFC.5F | TTCACCTTCTACATTTCACTATGTGC | 11 | 625 |
| NFC.5R | TTTATTCCTGCAGATCAATATTTCCC | 16 |  |
| NFC.6F | TGTGGAATCCTGATGCTCCTG | 15 | 637 |
| NFC.6R | CCTGCAGTGGGATGCTCAAT | 19 |  |
| NFC.7F | CACTGAAGCTGTTCTGGTTGC | 18 | 650 |
| NFC.7R | TGGTCCGTATTTGAAGTCCCAC | 21 |  |
| NFC.8F | TACAGGAATGGATCAACATGACTG | 21 | 588 |
| NFC.8R | AGAGGTCATCTCTCCTTGCCA | 23 |  |
| NFC.9F | CATCTAGGGCAAGCTAGCATTG | 22 | 496 |
| NFC.9R | GACCGTACAGTGCCTCAGTG | 26 |  |
| NFC.10F | GACTGGGTTATGGGAACATCAAAC | 24 | 595 |
| NFC.10R | TAACCAGAACTCGAGCTAGTTCAT | 28 |  |
| NFC.11F | GTCACAATGATGGGTGATCAAGG | 27 | 499 |
| NFC.11R | TCAGTAGGGAGTGGCAAGTT | 31 |  |
| NFC.12F | CAACTTGCCACTCCCTACTGA | 31 | 493 |
| NFC.12R | GGTCGTCTTCCAACAGCTTTAT | 35 |  |
| NFC.13F | CATAAGTGACGGCAATGTGCT | 34 | 601 |
| NFC.13R | AGTCAGCAGCCGCTCATGAT | 37 |  |
| NFC.14F | CCATACCGGGCCTAGCAATC | 37 | 696 |
| NFC.14R | CACAGAAGATTATAGGCAGCTGA | 39 |  |
| NFC.15F | ACTGTCACAGCCCGACTCTA | 38 | 417 |
| NFC.15R | ACTCTTTGTCGTTTGGCATCA | 40 |  |
| NFC.16F | GAGCCACACCTCACGTTAGAA | 39 | 697 |
| NFC.16R | GTCTCAAAACTTGCTTGGTCTCTT | 43 |  |
| NFC.17F | TCCGCTCTCCCTTAGAGCTT | 42 | 618 |
| NFC.17R | ACAGCTACCCAAAAGAGGGC | 47 |  |
| NFC.18F | GCACTTGAGAGTTGCTTAAAAGGAC | 46 | 558 |
| NFC.18R | CTTTCTATGTTTTAGGCTGCAGCGAC | 50 |  |
| NFC.19F | GGTACAGGCATCCTTCACCTGCTATTG | 49 | 676 |
| NFC.19R | ACAGTAAGAAGCAGCGCCTGG | 53 |  |
| NFC.20F | GACACAAAGGCTCCTAAAAGGC | 52 | 625 |
| NFC.20R | GTTTGGTATTGTGGTGGGGATT | 55 |  |
| NFC.21F | GCCAGTGTTGTGTTTCCCAAA | 54 | 691 |
| NFC.21R | CAGGAAGTGCAGCATTACAACAT | 58 |  |

*numbering based on isoform NM_001042492 containing alternative exon in GRD
