## Supplemental Table 5-6 for "Immortalization and Characterization of Schwann Cell Lines Derived from NF1 Associated Cutaneous Neurofibromas"

| Cell Line | Variant Classification | Variant Type | Genetic Change | Protein Change | dbSNP ID |
|---|---|---|---|---|---|
| 28cNF | Nonsense_Mutation | SNP | c.3233C>G | p.S1078* | rs2067122696 |
| i28cNF | Nonsense_Mutation | SNP | c.3158C>G | p.S1053* | rs1597717610 |
| i28cNF | Nonsense_Mutation | SNP | c.3233C>G | p.S1078* | rs2067122696 |
| cNF00.10a | Nonsense_Mutation | SNP | c.5861C>A | p.S1954* | |
| icNF00.10a | Nonsense_Mutation | SNP | c.5861C>A | p.S1954* | |
| cNF04.9a | Splice_Site | SNP | c.5268+2T>G | p.X1756_splice | rs1555533416 |
| cNF04.9a | Frame_Shift_Del | DEL | c.7391_7407del | p.T2464Kfs*13 | novel |
| icNF04.9a | Splice_Site | SNP | c.5268+2T>G | p.X1756_splice | rs1555533416 |
| icNF04.9a | Nonsense_Mutation | SNP | c.5503C>T | p.Q1835* | |
| icNF04.9a | Frame_Shift_Del | DEL | c.7391_7407del | p.T2464Kfs*13 | novel |
| cNF97.2a | Frame_Shift_Del | DEL | c.233del | p.N78Ifs*7 | rs1438566555 |
| cNF97.2a | Frame_Shift_Del | DEL | c.1929del | p.M643Ifs*45 | rs1567846742 |
| icNF97.2a | Frame_Shift_Del | DEL | c.233del | p.N78Ifs*7 | rs1438566555 |
| icNF97.2a | Frame_Shift_Del | DEL | c.1929del | p.M643Ifs*45 | rs1567846742 |
| cNF97.2b | Frame_Shift_Del | DEL | c.233del | p.N78Ifs*7 | rs1438566555 |
| cNF97.2b | Splice_Site | DEL | c.1392+2_1392+3del | p.X464_splice | novel |
| icNF97.2b | Frame_Shift_Del | DEL | c.233del | p.N78Ifs*7 | rs1438566555 |
| icNF97.2b | Splice_Site | DEL | c.1392+2_1392+3del | p.X464_splice | novel |
| cNF98.4c | Splice_Site | SNP | c.6704+1G>T | p.X2235_splice | rs1060500376 |
| icNF98.4c | Splice_Site | SNP | c.6704+1G>T | p.X2235_splice | rs1060500376 |
| cNF98.4d | Frame_Shift_Del | DEL | c.6316del | p.V2106Lfs*5 | |
| cNF98.4d | Splice_Site | SNP | c.6704+1G>T | p.X2235_splice | rs1060500376 |
| icNF98.4d | Frame_Shift_Del | DEL | c.6316del | p.V2106Lfs*5 | |
| icNF98.4d | Splice_Site | SNP | c.6704+1G>T | p.X2235_splice | rs1060500376 |

***Supplemental Table 5 - NF1 mutations detected by targeted sequencing are recapitulated using whole-genome sequencing.*** *Orange indicates primary cell cultures, while green indicates immortalized cell lines.*

| Primary cell line | Immortalized cell line | Jaccard index |
|---|---|---|
| 28cNF | i28cNF | 0.722 |
| cNF00.10a | icNF00.10a | 0.714 |
| cNF04.9a | icNF04.9a | 0.715 |
| cNF97.2a | icNF97.2a | 0.679 |
| cNF97.2b | icNF97.2b | 0.717 |
| cNF98.4c | icNF98.4c | 0.720 |
| cNF98.4d | icNF98.4d | 0.712 |

***Supplemental Table 6 - Jaccard indices of the WGS variant profiles identified in the paired primary and immortalized cell lines.***
